# Atlas-independent brain connectome analysis at voxel-level granularity: graph convolutional networks for etiology classification in newborns

**DOI:** 10.1101/2025.05.08.652584

**Authors:** Anna Speckert, Lukas Gianinazzi, Sepp Kollmorgen, Cornelia Hagmann, Patrice Grehten, Raimund Kottke, Giancarlo Natalucci, Beatrice Latal, Walter Knirsch, Ruth Tuura, Spina Bifida Study Group Zurich, Torsten Hoefler, Andras Jakab

## Abstract

Early identification of altered brain networks in neonates at risk for neurodevelopmental impairments is critical for timely intervention and improving outcomes. This study explores the potential of graph convolutional networks (GCNs) applied to structural brain connectomes at the voxel level granularity to classify neonatal connectomes by their underlying etiology: 51 children with congenital heart disease (CHD), 100 children born very preterm (PB), and 43 children with spina bifida aperta (SBA). Leveraging the flexibility of voxel-level parcellation, we captured fine-grained connectomic differences that improved classification performance (F1 = 0.78) compared to both atlas-based methods (F1 = 0.62) and a multilayer perceptron baseline model (F1 = 0.69). This approach enables subject-specific parcellation without the need for predefined anatomical templates, facilitating the analysis of diverse brain morphologies and age ranges. Attribution analysis using integrated gradients provided interpretable insights into etiology-specific connectomic patterns, highlighting regions of potential neurodevelopmental importance, such as the Rolandic operculum, inferior parietal lobule, and inferior frontal gyrus. Lateralized attribution patterns in PB reflected known neurodevelopmental alterations, underscoring the value of interpretable graph learning for understanding etiology-specific connectomic features. This work represents an important step toward atlas-independent connectome analysis, offering a novel framework for studying diverse neonatal populations and advancing our understanding of early brain development.

## 1. Introduction

Studying neonatal brain development is essential for understanding the origins of neurodevelopmental diseases and to identify individuals at highest risk for neurodevelopmental impairments. Many conditions associated with altered neurodevelopment, such as prematurity (McCormick et al., 2011), congenital heart disease (CHD) (Feldmann et al., 2021), and spina bifida aperta (SBA) (Schneider et al., 2021), are closely linked to disruptions in white matter development. These conditions are associated with an increased risk of cognitive and developmental impairments, highlighting the importance of studying early brain connectivity.

Disruptions in white matter can be studied using diffusion magnetic resonance imaging (dMRI) to map the brain’s structural networks, also known as the structural connectome. The neonatal period is a unique time to study the brain as it rapidly develops, forming and refining neural connections essential for development, which has been reflected by studies using dMRI (Dubois et al., 2014). This has given rise to many works aiming to reveal how the human brain connectome changes in pre- and postnatal development, and how congenital or acquired pathologies might affect connectomic structure (Bonthrone et al., 2022; Feldmann et al., 2020; Ní Bhroin et al., 2020; van den Heuvel et al., 2015). Methods for analyzing structural brain connectomes vary widely (Maier-Hein et al., 2017). Factors such as the choice of tractography algorithms (Nath et al., 2020; R. E. Smith et al., 2015b), atlases, region-of-interest granularity (Dadashkarimi et al., 2023), and thresholding (Tsai, 2018) significantly impact the reproducibility and biological validity of the results. Importantly, the majority of the dMRI based connectome analysis frameworks require the imaging data to be transformed to a standard space, or atlases, to reach anatomical correspondence between subjects. This presents a challenge in many scenarios, such as during fetal and neonatal brain development when images are difficult to standardize, in pathological conditions, or when conducting cross-etiologic or cross-age analyses (Hunt et al., 2019).

Once connectomes are obtained, several methods exist for the analysis of connectome development or between-group effects. Graph theory, a mathematical framework to analyze and quantify networks, has been widely applied to such data (Bullmore & Sporns, 2009; Papo et al., 2014; Rubinov & Sporns, 2010; van den Heuvel & Sporns, 2019; Zhao et al., 2019). It has helped uncover brain network disruptions in conditions such as Alzheimer’s disease (Yu et al., 2021), ADHD (Cao et al., 2014), Parkinson’s disease (Galantucci et al., 2017), schizophrenia (Fornito et al., 2012), and autism (Buckholtz & Meyer-Lindenberg, 2012) and revealed that connectome vulnerabilities can emerge as early as the prenatal stage (Batalle et al., 2012).

However, this approach has inherent constraints. It has been shown that the connectomic data (networks) need to meet certain boundary conditions to undergo edge-level of graph-theory based analysis. Among these conditions, a proportionally equal density of edges as well as an equal number and anatomical equivalence of nodes is required. Traditional graph theory analyses often abstract the connectome into a simplified network representation, which can fail to account for the spatial and anatomical relationships between brain regions that are integral to its topological organization, thereby limiting the ability to capture complex network characteristics (Rosenthal et al., 2018). Second, to apply traditional graph theory across subjects, brain networks must be of uniform size, which is typically achieved using brain atlases that parcellate the brain into predefined regions or nodes. These constraints pose challenges for comparing connectomes across populations or age ranges, where brain morphology varies significantly. Several solutions have been proposed to overcome these limitations. For example, Dadashkarimi et al. (2023) proposed using optimal transfer to generalize connectomic data. Others have aimed to decouple anatomical information from functional connectomic structure by aligning subjects in low dimensional spaces defined by connectome similarity (Kim et al., 2021; T. Xu et al., 2020). Interestingly, this challenge has gained momentum mostly in functional connectome analysis (i.e., fMRI-based), while machine learning-based methods are only beginning to emerge (Yang et al., 2025).

More recently, graph neural networks (GNNs), a family of deep learning models, have gained attention for their ability to learn powerful representations of brain network topologies (Cui et al., 2021). Among these, graph convolutional networks (GCNs) are particularly effective due to their simple inductive bias when aggregating and processing information from neighboring nodes, enabling them to learn hierarchical representations of graph structure (Kipf & Welling, 2016). By leveraging these representation-learning capabilities, GNNs have been applied to classify diseases such as HIV and bipolar disorder (Cui et al., 2021), Alzheimer’s disease (Subaramya et al., 2022), and autism spectrum disorder (Cao et al., 2021). They have also been used to distinguish between cognitive states (Li et al., 2021) and predict clinical outcomes such as cognitive performance (Wu et al., 2021). Another advantage of GNNs is their ability to handle graphs of varying sizes. This flexibility eliminates the need for predefined brain parcellations, such as atlas-based methods, and allows for the use of data-driven parcellations, such as voxel-based parcellation. Voxel-based parcellation generates graphs directly from the imaging data, capturing more granular information while avoiding biases introduced by predefined anatomical templates. It also facilitates the analysis of diverse populations and conditions by accommodating varying brain morphologies and providing uniformly sized nodes, making it particularly suited for comparative connectomics (de Reus & van den Heuvel, 2013).

Despite their powerful representation-learning capabilities, GNNs, like many deep learning models, are often criticized for their lack of interpretability, which limits their applicability in decision-critical contexts such as disease diagnosis (Cui et al., 2021). To address this, interpretability techniques like integrated gradients (Sundararajan et al., 2017a) have been developed to provide insights into model predictions. Integrated gradients attribute the model’s output to its input features, offering a transparent way to identify which input features contribute most to a classification decision.

We aim to explore the potential of GCNs for classifying neonatal brain connectomes by etiology. In our study, etiology classification serves as an important proof of concept: it demonstrates whether useful features are being learned and provides insight into connectome-level differences associated with conditions arising from various disruptions in white matter development in neonates. Specifically, we investigate whether an atlas-free connectome analysis approach can improve classification accuracy and offer a versatile framework for analyzing connectomes across diverse populations and age ranges. Additionally, to enhance the interpretability of our GCN-based framework, we explore using integrated gradients.

## 2. Methods

### 2.1 Study Populations

The data for this analysis were drawn from two sources: the publicly available Developing Human Connectome Project (dHCP), which includes neonatal data from healthy controls and preterms (Edwards et al., 2022), and three cohort studies conducted at the University Children’s Hospital Zurich.

#### 2.1.1 dHCP study population

For this study, we used the second neonatal release from the dHCP, accessed in January 2022. To achieve a balanced representation of gestational ages at the time of scanning, we randomly sampled the dataset to include approximately the same number of subjects per week within the range of corrected gestational ages between 25.6 and 42.3 weeks, thereby including both preterm and term-born infants. This also helped to balance the case numbers between the dHCP and the further cohorts used in the current study. The final dataset comprised MRI scans along with demographic information, including birth age, scan age, and sex, for a total of 120 subjects. After excluding five cases due to missing or low-quality diffusion MRI data, the final analysis included 115 participants (66 healthy controls and 49 preterm born infants).

The dHCP was approved by the National Research Ethics Committee in the United Kingdom and informed written consent was given by the parents of all participants.

### 2.1. University Children’s Hospital Zurich study population

Subject’s from three cohort studies conducted at the University Children’s Hospital Zurich were included in this analysis: “Heart and Brain” (HB) study (Meuwly et al., 2019), which enrolled neonates with complex congenital heart disease (CHD) and healthy controls; a prospective collection of spina bifida aperta (SBA) cases undergoing fetal repair surgery (Moehrlen et al., 2020); and a subset of subjects from “The Swiss EPO Neuroprotection Trial” (EPO) (O’Gorman et al., 2015), which focused on preterm infants born between 26 and 31 weeks of gestation. For this analysis, subjects were selected from these cohorts if high-quality T2-weighted and diffusion-weighted MR images were available (nControls = 42; nCHD = 54; nPB = 53; nSBA = 44). After excluding cases with image artifacts during processing, a total of 187 subjects from the University Children’s Hospital Zurich were included (nControls = 42; nCHD = 51; nPB = 51; nSBA = 43). Additional details about the cohort studies can be found in (Speckert, Payette, et al., 2024).

Informed written consent was obtained from all parents or caregivers for the use of their infants’ data in research. The ethical committee of the Canton of Zurich approved the collection and retrospective analysis of clinical data for the studies (approval numbers 2016-01019, 2021-01101, and 2022-0115). The clinical variables used in this work were derived from systematically and prospectively maintained data registries for each study. Data specific to spina bifida newborns were extracted from a REDCap™-based repository. The HB study received approval from the ethics committee of the Canton of Zurich, Switzerland (KEK StV-23/619/04), and was conducted in compliance with the principles outlined in the Declaration of Helsinki and Good Clinical Practice guidelines. For the EPO study, ethical approval was granted by the local ethical committee (KEK StV-36/04), and the study was approved by the Swiss drug surveillance unit (Swissmedic, 2005DR3179).

### 2.2 Magnetic resonance imaging (MRI) acquisition

#### 2.2.1 dHCP MRI acquisition

The T2-weighted images from the dHCP data were obtained using a Turbo Spin Echo (TSE) sequence, followed by processing with super-resolution reconstruction (Kuklisova-Murgasova et al., 2012) and motion correction (Cordero-Grande et al., 2018), resulting in a 3D T2-weighted brain volume with isotropic resolution of 0.5 × 0.5 × 0.5 mm³. For this study, the 3DT2 volumes were sourced from the open-access dHCP database. Details of the MRI acquisition parameters are described in Makropoulos et al. (2018).

The diffusion imaging protocol employed a spherically optimized scheme across four b-values (b0: 20; b = 400 s/mm²: 64; b = 1000 s/mm²: 88; b = 2600 s/mm²: 128), providing high angular resolution. Scans were acquired at a resolution of 1.5 × 1.5 mm in-plane with a slice thickness of 3 mm and imaging parameters of TR = 3800 ms, and TE = 90 ms. Further details about the diffusion MRI protocol are available in Bastiani et al. (2019).

#### 2.2.2 University Children’s Hospital Zurich MRI acquisition

The preterm infants were scanned on a 3.0 T GE HD.xt MRI scanner (GE Medical Systems) equipped with an 8-channel head coil. The scanner was later upgraded with new gradient coils (GE Signa MR750), but the same head coil continued to be used for neonatal MRI scans in the HB study, which included both CHD infants and healthy controls, as well as for the SBA infants. For the CHDs, healthy controls and preterm born infants, imaging was performed during natural sleep, whereas SBA infants were scanned under sedation with chloral hydrate.

The T2-weighted scanning protocol of the CHDs and healthy controls included a fast spin-echo sequence in axial, coronal and sagittal planes. The preterm and SBA cohort’s T2-weighted MRI was performed with a fast spin-echo sequence in axial, sagittal and coronal planes. After super-resolution reconstruction (Kuklisova-Murgasova et al., 2012), the resulting 3D T-weighted brain volumes had an isotropic resolution of 0.4 mm. Detailed parameters for each cohort are available in (Speckert, Payette, et al., 2024).

The diffusion imaging protocol for the preterm infants used a pulsed gradient spin-echo echo-planar imaging sequence, with TR = 9000 ms, TE = 77 ms, 21 diffusion encoding gradient directions at a b-value of 1000 s/mm2 and four b=0 images. The remaining cohorts acquired axial diffusion tensor imaging (DTI) data using a pulsed gradient spin-echo echo-planar imaging sequence (TR = 3950 ms, TE = 90.5 ms (for CHDs and healthy controls) and TE = 90 ms (for SBA), slice thickness=3 mm) with 35 diffusion encoding gradient directions at a b-value of 700 s/mm2 and four b=0 images. Further details about the diffusion MRI protocols can be found in the respective study protocols (Meuwly et al., 2019; Möhrlen et al., 2020; O’Gorman et al., 2015).

### 2.3 MRI preprocessing

The aim of the MRI processing was to generate whole-brain structural connectomes using anatomically constrained probabilistic tractography (ACT). To ensure comparability across datasets, the image processing steps were designed to be as consistent as possible. However, since the dHCP data were not raw but had undergone prior preprocessing, the following section outlines the initial preprocessing steps separately for the dHCP data and the University Children’s Hospital data.

#### 2.3.1 dHCP MRI preprocessing

After super-resolution reconstruction, the dHCP structural images were processed using the dHCP structural pipeline (Makropoulos et al., 2018), which segmented the T2-weighted images into 87 anatomical regions.

The diffusion data from the dHCP were pre-processed with the dHCP diffusion MRI pipeline (Bastiani et al., 2019), including denoising (Cordero-Grande et al., 2019) motion correction (Cordero-Grande et al., 2018) and eddy current distortion correction from FSL (Andersson & Sotiropoulos, 2016). Lastly, we applied B1 bias field correction from MRtrix3Tissue (https://3Tissue.github.io), an extension of MRtrix3 (Tournier et al., 2019), using Advanced Normalization Tools (ANTs) N4 (Tustison et al., 2010).

#### 2.3.2 University Children’s hospital Zurich MRI preprocessing

For the University Children’s hospital Zurich data, the same image processing pipeline has been used for all the cohorts as described in (Speckert, Payette, et al., 2024).

The super-resolution reconstructed T2-weighted images were bias field corrected using ANTs N4 bias field correction (Tustison et al., 2010) and subsequently segmented into seven tissue types: cortical gray matter, deep gray matter, white matter, external cerebrospinal fluid (CSF), ventricles, brainstem, and cerebellum. Segmentation was performed using an in-house algorithm built on a U-Net architecture (Payette et al., 2021), trained on images from both healthy appearing brains and spina bifida cases.

The diffusion data was preprocessed using several steps. First, denoising was performed with the deep learning algorithm Patch2Self from the DIPY package in Python (Fadnavis et al., 2020). Next, Gibbs ringing artifacts were suppressed using the method described by Kellner et al. (2016), implemented in MRtrix3Tissue, to eliminate high-frequency oscillations. Finally, motion and distortion corrections were applied using slice-to-volume eddy correction with CUDA 8.0 in FSL version 6.00 (S. M. Smith et al., 2004). Lastly, B1 bias field correction was performed using ANTs N4 (Tustison et al., 2010) from MRtrix3Tissue.

### 2.4 Network construction: for all datasets

The segmented T2-weighted images were transformed into a 5-tissue type to cohere to MRtrix3’s requirement to perform ACT.

Brain parcellation was performed using two different approaches. The first approach utilized the Edinburgh Neonatal Atlas (ENA33), derived from the Desikan-Killiany atlas (Desikan et al., 2006) for neonatal brains (Blesa et al., 2016). The atlas template was registered to each subject’s T2-weighted image using the SyN method in ANTs (Avants et al., 2014). For spina bifida cases, an SBA specific neonatal atlas (OSBA) was used (Speckert, Ji, et al., 2024). The registered atlas was then aligned with the DTI space using the TRSAA + SyNOnly methods (Avants et al., 2014). The second approach involved voxel-wise parcellation. Using tissue segmentation, the gray matter of each subject was divided into individual voxels, enabling subject-specific parcellation without relying on predefined anatomical regions. However, to enable comparisons and analyses across the two parcellation methods, a mapping was created to assign each voxel node to its corresponding atlas label based on spatial overlap. This mapping ensured that voxel-based nodes could be grouped or interpreted in the context of the atlas regions, providing a consistent framework for integrating the two approaches.

Due to incomplete brain coverage in 40 DTI scans from the Children’s University Hospital Zurich, cerebellar atlas regions were excluded from the analysis for all subjects. Additionally, the lateral ventricles and corpus callosum were removed, as they do not contribute meaningfully as nodes in the structural brain connectome. This resulted in a modified atlas consisting of 94 regions. Further details are provided in (Speckert, Payette, et al., 2024).

Tissue response functions for white matter and CSF were estimated and averaged across 15 randomly selected subjects per study population using MRtrix3Tissue (Dhollander et al., 2019). Based on these averages, tissue and free water orientation distribution functions (ODFs) were calculated with the multi-shell, multi-tissue constrained spherical deconvolution algorithm in MRtrix3Tissue, treating b-value 0 as a shell. The ODFs were then normalized per subject to enable quantitative density measurements (Dhollander et al., 2021; Tournier et al., 2019).

ACT (R. E. Smith et al., 2012; Tournier et al., 2010) with dynamic seeding (R. E. Smith et al., 2015a) was used to generate 10 million streamlines. SIFT2 (R. E. Smith et al., 2015a; Tournier et al., 2019) was applied to align streamline densities within the white matter with fiber densities estimated by the spherical deconvolution model. To achieve inter-subject normalization of connection densities, the SIFT2 proportionality coefficient (µ) was calculated for each subject.

Following these preprocessing steps, structural connectomes were constructed using both atlas-based and voxel-wise parcellation approaches. For the atlas-based parcellation, the 94 regions from the modified ENA33 atlas served as nodes, with edges representing the connection strength, quantified as the sum of weighted streamlines (SIFT2 * µ) connecting each region. This resulted in a fixed 94×94 connectivity matrix for each subject. For the voxel-wise parcellation, nodes were defined as individual gray matter voxels, producing connectomes of varying sizes due to subject-specific differences in the number of gray matter voxels. Edges, representing the connection strength, were similarly calculated as the sum of weighted streamlines (SIFT2 * µ) connecting the voxels. More details on the resulting connectivity matrices can be found in Table 2.

### 2.5 Graph convolution network

To classify neonatal structural connectomes, we utilized a GCN. In the following, the model and training specifications are described.

#### 2.5.1 Model

##### 2.5.1.1 Features

We used the following seven input node features:

- The physical spatial coordinates (𝑥,𝑦, 𝑧) of the node in real-world scanner space, derived from the MRI scan and expressed in millimeters relative to the scanner’s coordinate system.
- The total connection strength, i.e., the sum of all edge weights.
- The directed connection strength provides a fixed-size representation of local connectivity for each vertex, capturing directional patterns in the connectome. This feature efficiently captures important connectivity patterns without needing to store every individual connection. Specifically, the directed connection strengths (1) compress complex neighborhood information into just three values, (2) retain information about which spatial directions have stronger connections, and (3) allow the model to understand spatial orientation without requiring atlas alignment. The directed connection strength for each coordinate ordering (𝑥,𝑦, 𝑧), (𝑦, 𝑥, 𝑧), and (𝑧, 𝑥,𝑦), is defined as follows: For a given coordinate 𝑖, an edge is included if the 𝑖-coordinate of the target node is increasing, or if it is equal and the subsequent coordinates satisfy the same condition recursively. As an example, consider two nodes u and 𝑣 with coordinates (0,1,2) and (2,1,0), respectively, connected by an edge of strength w. The directed connection strength is determined by evaluating coordinate orderings. In the (x, y, z) ordering, the first coordinate increases (0 → 2), so the edge is included with strength w. Similarly, in the (y, x, z) ordering, the first coordinates are equal (1 → 1), but the second coordinate increases (0 → 2), again including the edge with strength w. However, in the (z, x, y) ordering, the first coordinate decreases (2 → 0), excluding the edge with strength 0. Thus, this edge is included in the first two orderings but excluded in the third.

Each input feature was standardized to have mean 0 and standard deviation 1. For the directed connection strength, the mean and standard deviation were computed jointly across all three coordinate orderings. To simplify the approach and speed up training, edge features were not used. Instead, edge weights were encoded directly into the node features as described above.

##### 2.5.1.2 GCN architecture

Our network is based on a graph convolution network (GCN) (Kipf & Welling, 2016). We use the ReLU activation function denoted 𝜎 (Agarap, 2018) and, like the GIN layer (K. Xu et al., 2019), avoid degree normalization for more powerful aggregation. Given a node 𝑣, let 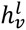 denote its representation at layer 𝑙 and 𝑊_*l*_ the weight matrix. Each layer is updated as:

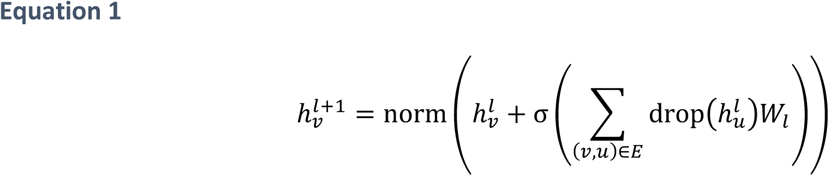

Here, drop(·) applies dropout (Srivastava et al., 2014) with probability 0.2 and norm(·) denotes graph instance normalization (Cai et al., 2021). The input features are projected into a 48-dimensional hidden space using a linear layer with ReLU activation. For the readout, we apply max-pooling over all nodes followed by two fully connected layers (Murtagh, 1991) with skip connections (He et al., 2016) and dropout. We used one convolutional layer in our GCN. As a non-convolutional baseline, we remove the graph convolution, resulting in a multi-layer perceptron (MLP).

##### 2.5.1.3 Graph Pruning

The degree distribution in voxel connectomes has a very long tail (see Figure 1). This can cause aggregation imbalances in Equation 1, where some few nodes with many neighbors dominate the sum. To address this, we limit each node to its top 256 edges (based on connection strength). The cutoff value of 256 edges was chosen based on initial experiments using a different fold seed and model initialization seed. This pruning significantly improved the learning outcomes (see <u>3.2</u> <u>Classification results</u> for details).

**Figure 1.**
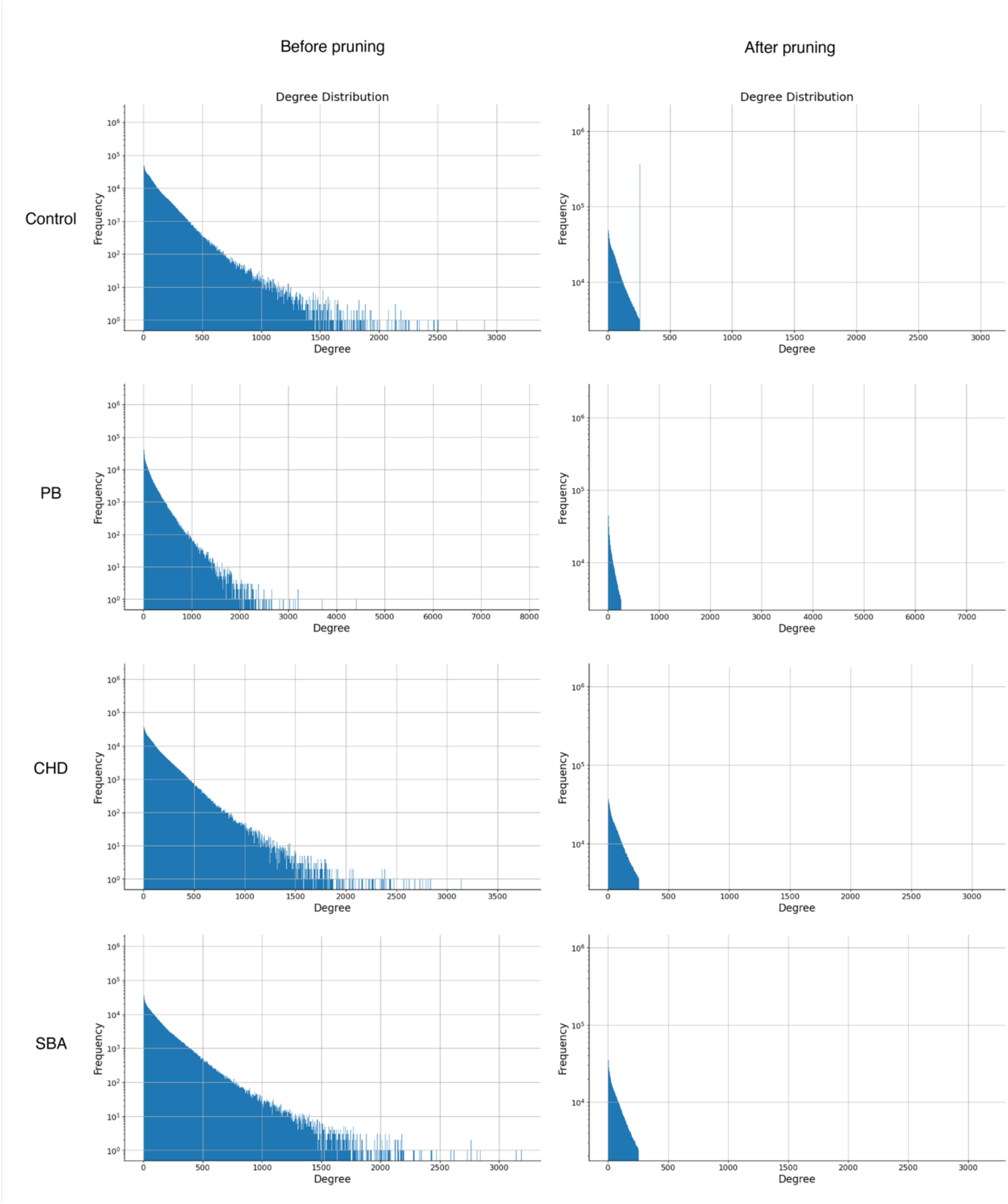
**Nodal degree distributions before and after pruning** *Note.* Log-scaled nodal degree distributions for voxel-based connectomes across etiologies, shown before (left column) and after pruning (right column). PB = premature birth; CHD = congenital heart disease; SBA = spina bifida aperta.

#### 2.5.2 Training

We trained using 5-fold cross-validation with three random seeds per fold (15 total runs). For optimization, we used AdamW (Loshchilov & Hutter, 2017) with a one-cycle learning rate scheduled (L. N. Smith & Topin, 2017) for 400 epochs and a batch size of 16. The learning rate was selected via a learning rate test (L. N. Smith, 2017). For the embedding visualizations and attribution analysis described in <u>2.7 Visualizations</u>, we trained a model on the full data using the same hyperparameters and a fresh random seed. Training took less than 8 hours per run on an Intel® Xeon® Gold 6154 Processor with 384 GB memory.

To mitigate class imbalance, we used stratified sampling. That is, we sample training examples inversely proportional to their class frequency. This ensured each batch contains, in expectation, an equal number of samples from each etiology. Cross-entropy loss was used for optimization.

### 2.6 Attribution Using Integrated Gradients

To understand which nodes contribute most to a given network decision, we use Integrated Gradients (IG) as an attribution method (Sundararajan et al., 2017b). IG quantifies the contribution of an input feature at a node to shifting the network output relative to a baseline output. This method has shown effectiveness in various graph learning tasks (McCloskey et al., 2019; Sanchez-Lengeling et al., 2020).

Let 𝐹: 𝑅^+×-^ → [0,1] be the network function mapping input features 𝑋 ∈ 𝑅^+×-^ (for 𝑁 nodes and 𝐷 features per node) to an output. A positive attribution value indicates that the feature increases the output relative to the baseline, while a negative value indicates a decrease.

When the baseline input 𝑋_0_ is the all-zero matrix, the (IG) for the 𝑖-th feature at node 𝑣 are defined as:

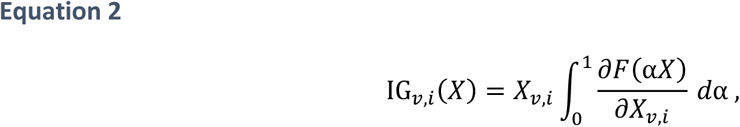

where α ∈ [0,1] interpolates between the baseline 𝑿_0_ and the actual input 𝑿. The term 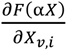 represents the partial derivative of 𝐹 with respect to the 𝑖-th feature at node 𝑣.

For the etiology classification problem, the all-zero matrix is used as the baseline. Each of the four output logits corresponds to a separate attribution task. To compute the total attribution for a node 𝑣, we sum the IG contributions across all features 𝑖:

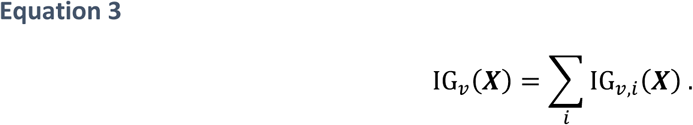

### 2.7 Visualizations

To explore and visualize the embeddings learned by the GCN, we applied Uniform Manifold Approximation and Projection (UMAP) (McInnes et al., 2018) for dimensionality reduction and explored using the *dataspace* software from the high-dimensional data science platform from the University of Zurich (Kollmorgen et al., 2020). Embeddings refer to a representation of data in a lower-dimensional vector space, where high-dimensional data, such as graph nodes, are mapped into a continuous space, with similar items positioned closer together. The UMAP algorithm was configured with 10 nearest neighbors, 1000 epochs, and Euclidean distance as the metric.

To investigate the importance of brain regions contributing to etiology classification, we used attribution values derived from IG to compute the inverse percentiles of mean node attribution for each region of interest (ROI). Voxel-based attributions were first mapped to brain regions defined by the ENA33 atlas (see <u>2.4 Network construction: for all datasets</u>), associating each voxel with its corresponding ROI. For each ROI, the mean node attribution was calculated by averaging the attribution values of all voxel nodes within the region across all subjects. To enhance interpretability and improve visual differentiation, these ROI-level mean attributions were transformed into inverse percentiles. Finally, the inverse percentiles supporting the classification of a specific etiology (e.g., Control) were examined separately across all four etiologies, enabling the exploration of attribution patterns for one etiology while considering data from all etiological groups.

To present the results, we visualized the attributions spatially and regionally. Spatial distributions were displayed along anatomical axes (e.g., coronal and axial slicing) using the XYZ coordinates of the ROIs, while regional distributions highlighted differences in attribution across specific brain regions. These analyses, including ROI aggregation, attribution averaging and visualization, were performed using the dspace software in MATLAB (version 2022a). The code can be found in the Supplementary material (2. MATLAB Code).

## 3. Results

### 3.1 Descriptive statistics

The analysis comprised a total of 302 subjects (187 from the University Children’s Hospital and 115 from the dHCP dataset). We revealed significant differences across the four groups in gestation age (GA) at birth (p < 0.0001, Kruskal-Wallis test H(3) = 236.93), postmenstrual (PMA) age at scan (p < 0.0001, Kruskal-Wallis test H(3) = 102.05) and sex (Chi-square = 43.88, df = 3, p < 0.0001).

Post-hoc Dunn tests with Bonferroni correction for GA at birth indicated that only the CHD and Control groups did not significantly differ (p = 1), while significant differences were observed between the remaining pairs of groups (Control vs. PB p < 0.0001, Control vs. SBA p < 0.0001, CHD vs. PB p < 0.0001, CHD vs. SBA p < 0.0001, PB vs. SBA p = 0.0007). Regarding PMA at scan, the post-hoc Dunn test revealed that only the SBA and PB groups did not significantly differ (p = 0.32), while significant differences were observed between the remaining pairs of groups (Control vs. CHD p = 0.005, Control vs. PB p < 0.0001, Control vs. SBA p < 0.0001, CHD vs. PB p = 0.0005, CHD vs. SBA p < 0.0001). For sex differences, the pairwise proportionality tests with Bonferroni correction revealed a significant difference only between the CHD and SBA group (p = 0.014). Further details about the sample characteristics can be found in Table 1.

**Table 1.** Sample characteristics of the study populations.

| Sample characteristic | Control<br>n = 108 | PB<br>n = 100 | CHD<br>n = 51 | SBA<br>n = 43 | p-value |
| --- | --- | --- | --- | --- | --- |
| GA at birth<br>(Mdn[IQR]) | 39.8<br>(37.8-41.8) | 30.9<br>(27.5-34.3) | 39.4<br>(37.4-41.5) | 36.0<br>(32.9–39.1) | p < 0.0001<sup>a</sup> |
| PMA at scan<br>(Mdn[IQR]) | 41.9<br>(38.9-44.9) | 37.4<br>(31.1-43.7) | 40.3<br>(38.2-42.5) | 38.0<br>(36.7-39.3) | p < 0.0001<sup>a</sup> |
| Male sex (N[%]) | 51 (47.2%) | 57 (57%) | 36 (70.6%) | 16 (37.2%) | p < 0.0001<sup>b</sup> |

*Note.* N = 302. Mdn = Median. IQR = Interquartile range. <sup>a</sup>Kruskal-Wallis test, <sup>b</sup>Chi-square test. PB = premature birth, CHD = congenital heart disease, SBA = spina bifida aperta, GA = gestational age, PMA = postmenstrual age.

**Table 2.** Characteristics of voxel-based connectomes per etiology.

|  |  | Control | PB | CHD | SBA |
| --- | --- | --- | --- | --- | --- |
| Number of nodes | Mean | 109k | 83k | 99k | 90k |
|  | SD | 25k | 40k | 19k | 14k |
|  | Min | 64k | 21k | 57k | 59k |
|  | Max | 190k | 175k | 134k | 121k |
| Number of edges before pruning | Mean | 9'632k | 9'176k | 9'461k | 8'193k |
|  | SD | 1'032k | 1'187k | 1'107k | 997k |
|  | Min | 5'553k | 5'859k | 5'884k | 5'772k |
|  | Max | 11'554k | 11'164k | 11'201k | 9'905k |
| Number of edges after pruning | Mean | 7'997k | 6'442k | 7'715k | 6'677k |
|  | SD | 922k | 1'314k | 914k | 829k |
|  | Min | 4'978k | 3'327k | 4'754k | 4'861k |
|  | Max | 10'446k | 9'158k | 9'106k | 8'143k |
| Nodal<br>degree<br>distribution | Median | 27 | 24 | 40 | 36 |
|  | Q1 | 1 | 1 | 1 | 1 |
|  | Q3 | 129 | 142 | 132 | 122 |

*Note.* Values expressed with 'k' are in thousands (e.g., '109k' represents 109,000). Other numbers are shown in their original scale. Q1 = first quartile; Q3 = third quartile; SD = standard deviation; PB = premature birth; CHD = congenital heart disease; SBA = spina bifida aperta.

To provide an overview of the connectomes generated using voxel-level parcellation, Table 2 summarizes key characteristics, complemented by histograms in Figure 1 for further visualization. Across all groups, degree distributions were right-skewed, with a long tail indicating a small subset of highly connected nodes. Pruning reduced the spread but preserved the overall shape of the distributions, with most node degrees falling below 500 across all groups. Notably, the total number of nodes and edges varied across groups, with healthy control and CHD connectomes generally showing higher values than PB and SBA.

### 3.2 Classification results

We evaluate our model using F1-score, precision, and recall. To handle class imbalance, we compute the scores using macro-averaging, where the metric is calculated independently for each class and averaged equally across all classes. Note that this means that the macro-average F1 score is not a function of the macro-average precision and recall. For each run, we aggregate the macro-averaged scores. We then report the mean and standard deviation of these scores across all 15 runs to ensure robust performance evaluation (see Table 3).

**Table 3.** Comparison of etiology classification performances.

|  | F1 | Precision | Recall |
| --- | --- | --- | --- |
| **atlas parcellation with MLP** | 0.41 | 0.44 | 0.61 (0.04) |
| mean (SD) | (0.05) | (0.07) |  |
| **atlas parcellation with GCN** | 0.62 | 0.63 | 0.75 (0.04) |
| mean (SD) | (0.07) | (0.07) |  |
| **voxel parcellation with MLP** | 0.69 | 0.71 | 0.8 |
| mean (SD) | (0.05) | (0.05) | (0.03) |
| **voxel parcellation with GCN** | 0.7 | 0.72 | 0.8 |
| mean (SD) | (0.08) | (0.09) | (0.05) |
| **voxel parcellation with GCN + pruning** | **0.78** | **0.79** | **0.86** |
| mean (SD) | (0.05) | (0.07) | (0.03) |

Note. MLP = multilayer perceptron; GCN = graph convolutional network; SD = standard deviation.

#### 3.2.1 Embedding visualization

**Error! Reference source not found.** shows the UMAP visualizations of the graph embeddings across different layers of the model.

In the supplementary materials, additional UMAP visualizations are provided. These visualizations show the connectome embeddings colored according to scanner ID, sex, age at birth, and age at scan and reveal that the GCN embeddings capture etiology-specific patterns while minimizing the influence of the confounding factors (Supplementary Figure 1). Furthermore, four subject-specific examples (one per etiology) are included to demonstrate how individual embeddings relate to the overall clustering (Supplementary Figure 2).

### 3.3 Integrated Gradients

Figure 3 shows the mean node attribution per etiology, illustrating the variability of attribution contributions for each group.

**Figure 2.**
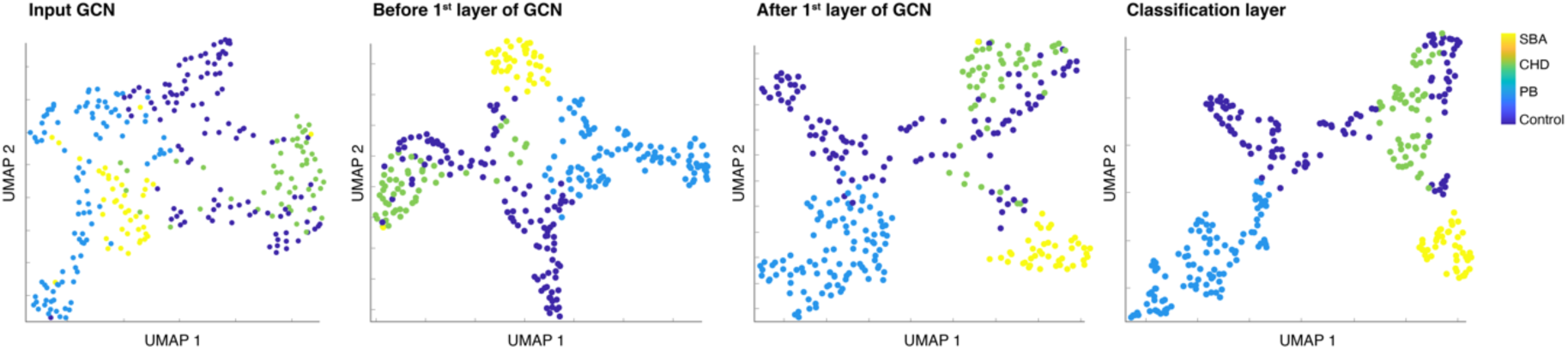
**UMAP visualization of the connectome embedding** *Note.* Embeddings colored by etiology UMAP configured with 10 nearest neighbors, 1000 epochs, Euclidean distance. PB = premature birth; CHD = congenital heart disease; SBA = spina bifida aperta.

**Figure 3.**
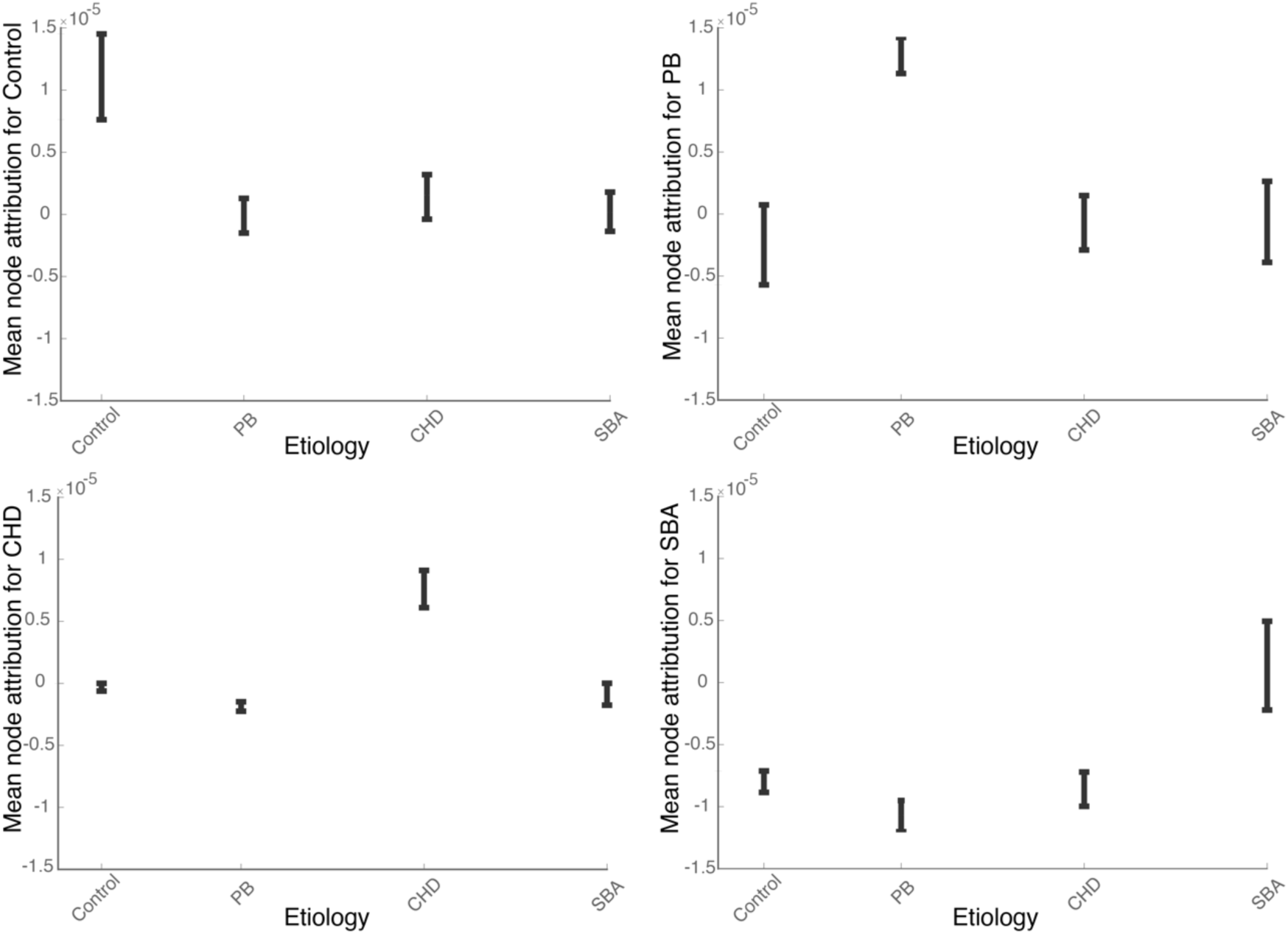
**Mean node attribution per etiology** *Note.* Depicted are the 95% bootstrapped confidence intervals, which indicate the variability of attribution estimates across subjects.

While Figure 3 provides an overview of the mean node attributions per etiology, the following figures focus on their spatial and regional distributions. To visualize how specific brain regions contribute to the classification of each etiology, we began by averaging the voxel-based attribution values within each ENA33-defined ROI for each subject, separately for the attribution values corresponding to each etiology. This process was repeated for all four etiologies. We then calculated the inverse percentiles of these averaged attributions to enhance subtle differences, making the visualizations more interpretable. Finally, we plotted these inverse percentiles using the ENA33-ROI coordinates, creating separate visualizations for each etiology based on their specific attribution values. These brain-like visualizations highlight how specific regions contribute to the classification of each etiology: Figure 4 for healthy Controls, Figure 5 for PB, Figure 6 for CHD, and Figure 7 for SBA. For each etiology, the five ROIs with the highest attribution values were identified and are listed in descending order of attribution. These regions include right occipital superior gyrus, right frontal superior medial gyrus, left occipital superior gyrus, right calcarine sulcus and right olfactory sulcus for the healthy controls; the right postcentral gyrus, right frontal middle gyrus, right parietal inferior gyrus, right Rolandic operculum, and right frontal inferior triangularis for PB; the left Heschl’ gyrus, left parietal inferior gyrus, left Rolandic operculum, left frontal inferior gyrus opercularis, and left temporal superior gyrus for CHD; and the left lingual gyrus, right temporal superior gyrus, left thalamus, left paracentral lobule, and left pallidum for SBA.

**Figure 4.**
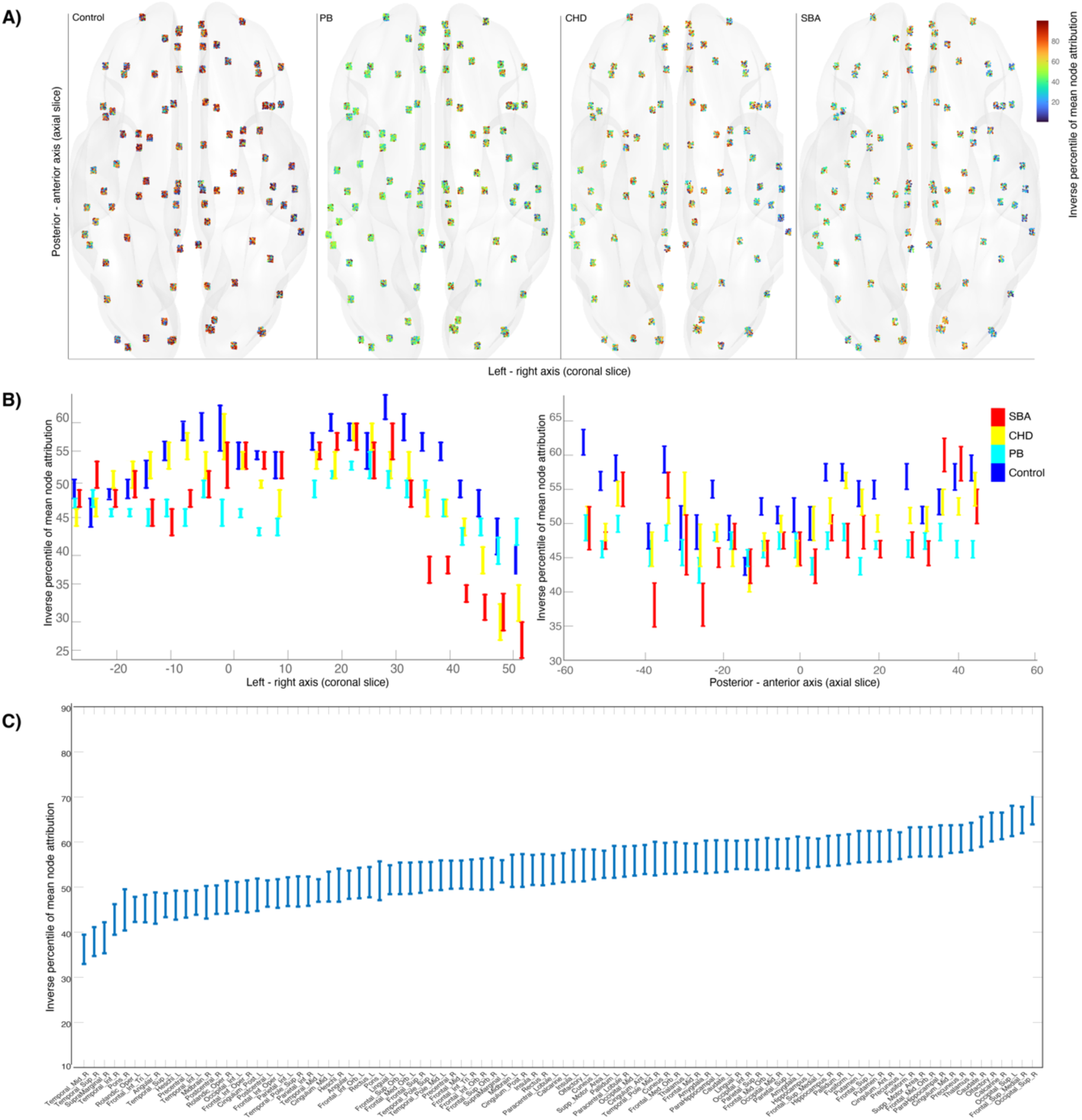
**Spatial and regional distribution of attribution supporting control classification** *Note.* (A) Visualizes the spatial distribution of attributions across the brain in the posterior-anterior (y-axis) and left-right (x-axis) anatomical axes, colored by attribution values. (B) Shows the inverse percentiles of mean node attributions along the left-right (coronal) and posterior-anterior (axial) axes with 95% confidence intervals for the four etiologies. (C) Displays the regional distribution of attributions averaged per ROI using the ENA33 atlas, with red circles indicating the five most salient ROIs.

**Figure 5.**
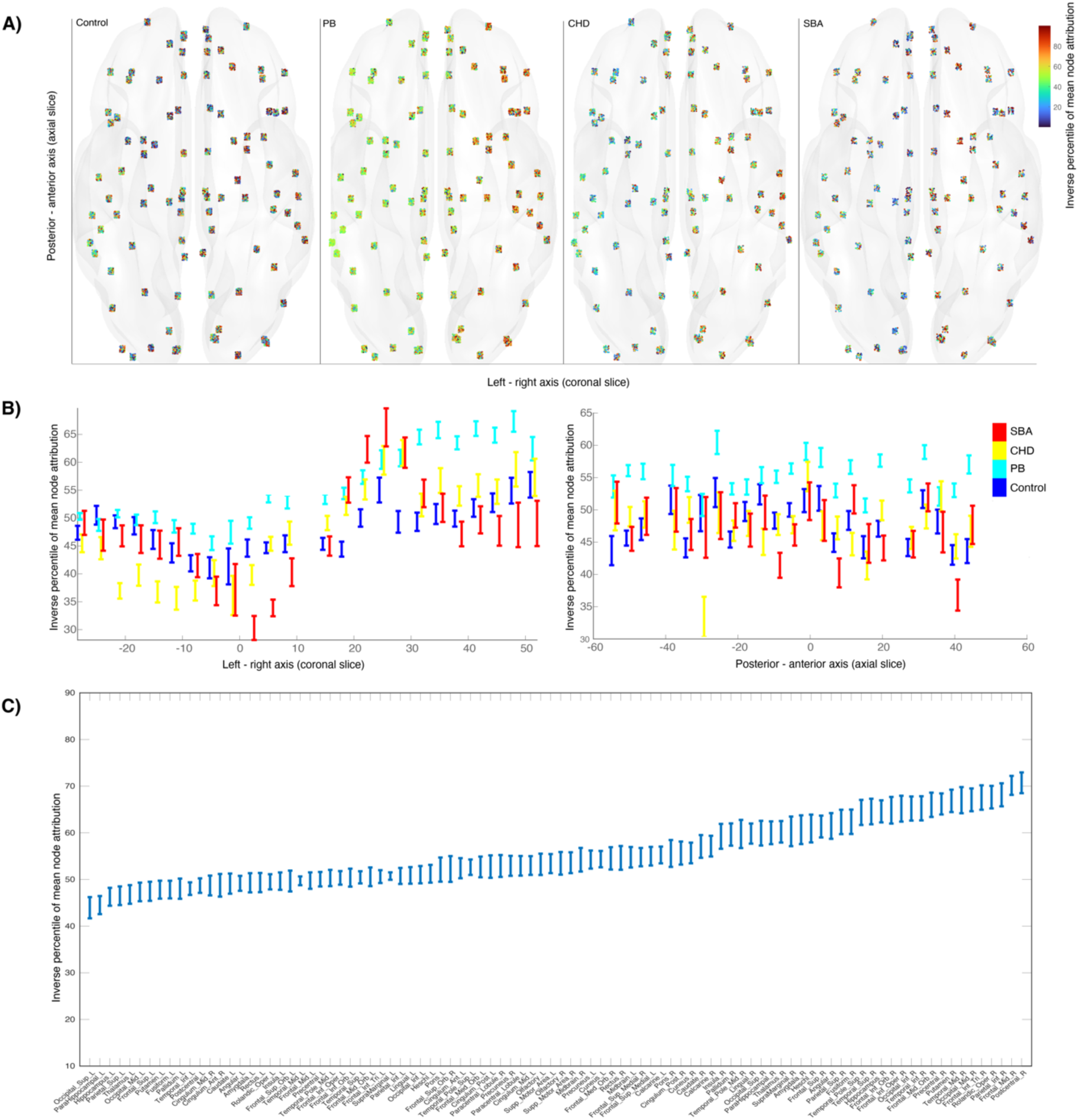
**Spatial and regional distribution of attribution supporting PB classificatio** *Note.* (A) Visualizes the spatial distribution of attributions across the brain in the posterior-anterior (y-axis) and left-right (x-axis) anatomical axes, colored by attribution values. (B) Shows the inverse percentiles of mean node attributions along the left-right (coronal) and posterior-anterior (axial) axes with 95% confidence intervals for the four etiologies. (C) Displays the regional distribution of attributions averaged per ROI using the ENA33 atlas, with red circles indicating the five most salient ROIs.

**Figure 6.**
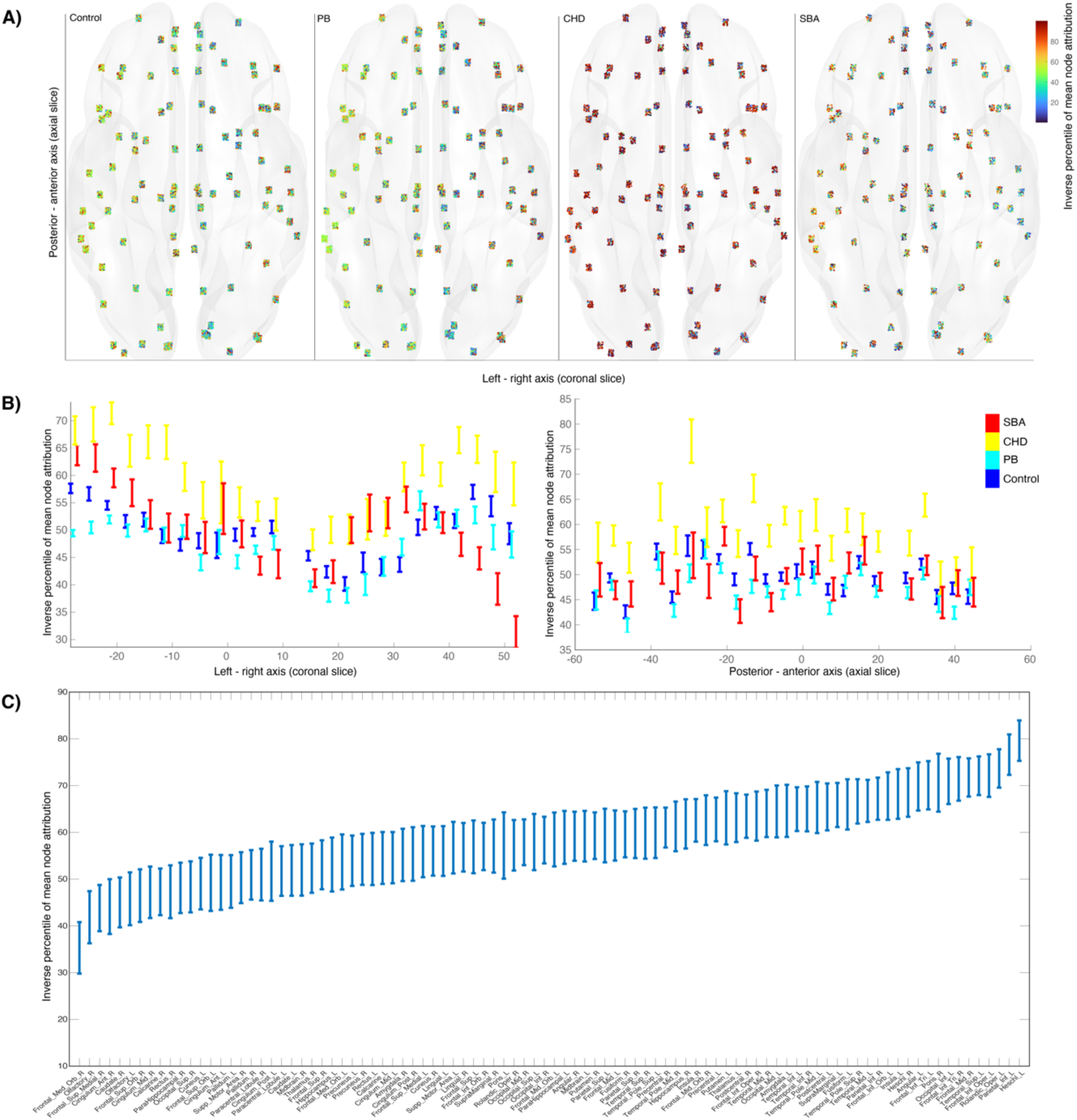
**Spatial and regional distribution of attribution supporting CHD classification** *Note.* (A) Visualizes the spatial distribution of attributions across the brain in the posterior-anterior (y-axis) and left-right (x-axis) anatomical axes, colored by attribution values. (B) Shows the inverse percentiles of mean node attributions along the left-right (coronal) and posterior-anterior (axial) axes with 95% confidence intervals for the four etiologies. (C) Displays the regional distribution of attributions averaged per ROI using the ENA33 atlas, with red circles indicating the five most salient ROIs.

**Figure 7.**
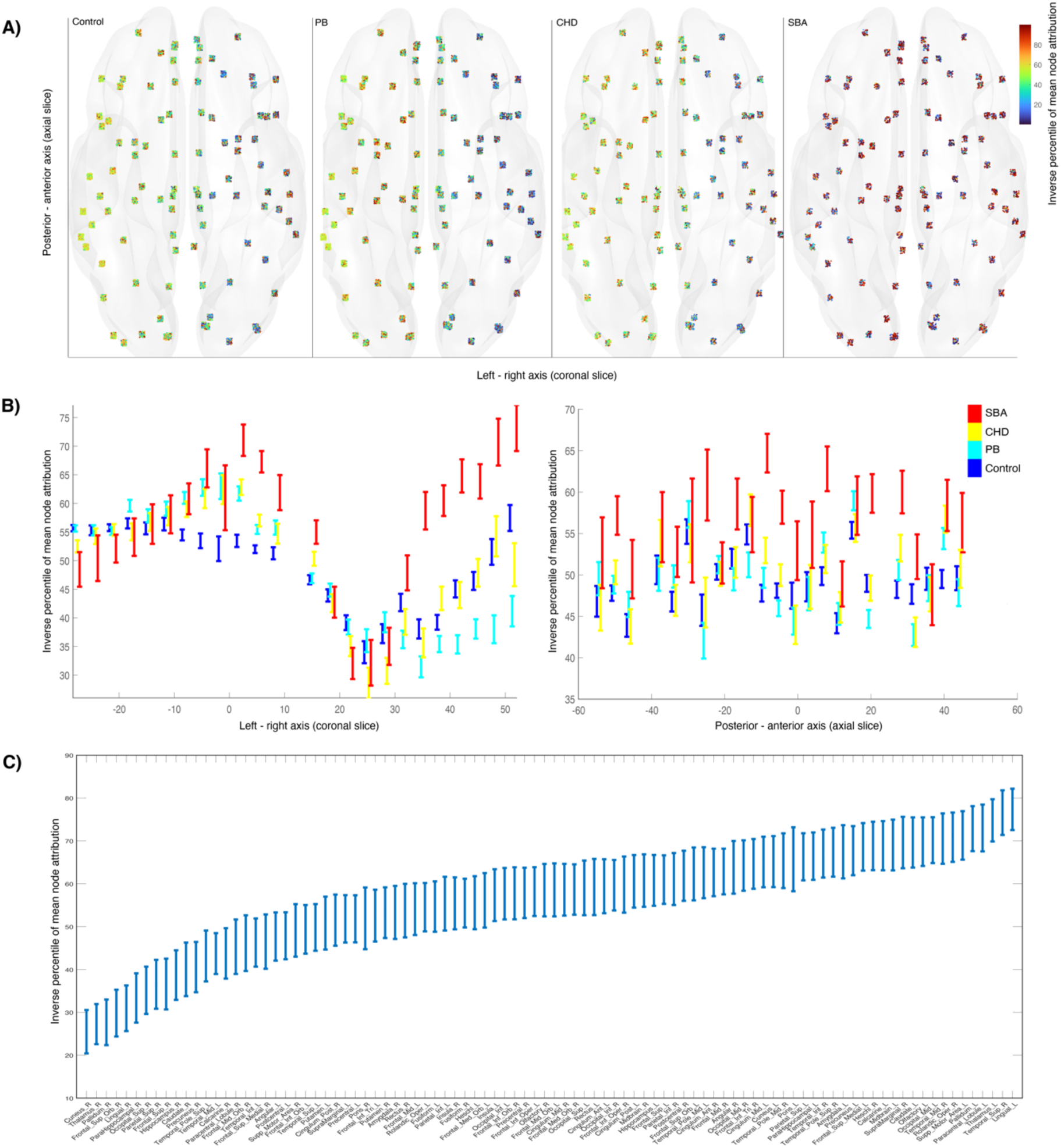
**Spatial and regional distribution of attribution supporting SBA classification** *Note.* (A) Visualizes the spatial distribution of attributions across the brain in the posterior-anterior (y-axis) and left-right (x-axis) anatomical axes, colored by attribution values. (B) Shows the inverse percentiles of mean node attributions along the left-right (coronal) and posterior-anterior (axial) axes with 95% confidence intervals for the four etiologies. (C) Displays the regional distribution of attributions averaged per ROI using the ENA33 atlas, with red circles indicating the five most salient ROIs.

## 4. Discussion

In this study, we demonstrated the utility of GCNs for classifying neonatal connectomes into distinct etiologies and examined the associated attribution patterns. Our results showed that voxel-based parcellation significantly improved classification performance compared to atlas-based parcellation, highlighting the advantage of higher spatial resolution in capturing connectomic differences. Furthermore, the GCN consistently outperformed the MLP across all parcellation strategies, underscoring the benefit of incorporating graph convolutional layers for learning topological features of the connectome.

In terms of IG analysis, we observed distinct patterns across etiologies. Notably, PB exhibited highly lateralized attributions distributions, whereas CHD’s attributions distribution was more similar to that of healthy controls, with a tendency towards higher importance in the left and right lateral hemisphere. Interestingly, SBA showed a trend within each hemisphere, with higher attribution values favoring the right side: the medial part of the left hemisphere and the lateral part of the right hemisphere displayed the highest attribution values. These findings emphasize the etiology-specific nature of attribution distributions and provide insights into the regional contributions underlying the model’s predictions. Moreover, they highlight the potential of attribution methods for revealing etiology-specific connectomic alterations that may not be apparent through traditional analyses.

### 4.1 Classification

Our classification results (Table 3) demonstrate that voxel-based parcellation significantly improves performance compared to atlas-based parcellation. This finding suggests that the higher spatial resolution provided by voxel-based connectomes not only introduces additional complexity and noise but also enhances the model’s ability to capture finer-grained connectomic details. Voxel-based connectomes have also been explored in other studies, such as Zalesky et al. (2012), which investigated the effects of long-term cannabis use on axonal fibre connectivity and demonstrated the potential of this approach to reveal subtle connectivity alterations. Moreover, the GCN consistently outperformed the MLP across both parcellation strategies, highlighting the advantage of graph convolutional layers in extracting topological features. The best performance was achieved using voxel-based parcellation with GCN, yielding an F1-score of 0.78. To our knowledge, this is the first study to apply GCNs for neonatal connectome-based etiology classification, demonstrating the utility of graph-based machine learning for connectome studies.

When examining the UMAP visualizations of the different GCN stages (Figure 2), the progressive separation of the etiologies becomes apparent. Healthy controls and CHD are the two groups that exhibit the least separation, with the controls showing a notable spread. This variability among controls could be attributed to their inclusion from two different datasets: the dHCP dataset and the HB study conducted at the University Children’s Hospital Zurich. While all dHCP controls were scanned using the same MRI scanner without changes, the HB study controls were scanned on the same scanner model, but a hardware upgrade introduced potential variability in the imaging parameters. Supplementary Figure 3 (top row) illustrates the embeddings of only the controls, colored by ScannerID. The dHCP scanner cohort appears to be clearly separated, whereas the scanner software upgrade did not substantially influence the graph learning. A similar trend is observed in the PB cohort embeddings with respect to ScannerID (Supplementary Figure 3, bottom row). Initially, the dHCP PBs are distinctly separated from the PBs from the University Children’s Hospital Zurich. However, as the embeddings progress through the GCN layers, these groups converge, indicating the model learns etiology-specific patterns rather than being confounded by ScannerID. Also, when examining the embeddings colored by potential confounding variables, a consistent pattern emerges. Age at birth shows stratification that is inherently linked to the etiologies and therefore persists throughout the layers (Supplementary Figure 1, third row). Scanning age initially introduces some separation, particularly among PBs, but its influence diminishes as the model progresses, reflecting its decreasing role in the GCN’s learning (Supplementary Figure 1, fourth row). Finally, sex does not show distinct patterns, corroborating the non-confounding nature of sex in the etiology classification (Supplementary Figure 1, second row). These findings underscore the GCN’s ability to focus on meaningful connectomic features for classification. Additionally, we tested correlation distance for UMAP dimensionality reduction, which produced qualitatively similar results to Euclidean distance. This consistency suggests that the choice of distance metric did not substantially influence the visualization of embeddings, further reinforcing the robustness of our observations.

### 4.2 Integrated gradients

To enhance the interpretability of the graph learning model, we used IG to assign attributions to specific ROIs. IG was selected as our attribution method due to its robustness and scalability, making it well-suited for analyzing voxel-based connectomes with varying graph sizes (Sanchez-Lengeling et al., 2020). Unlike the method proposed by Cui et al. (2022) which employs a dense matrix-based learned mask for edge-level attributions, IG provides node-level attributions through an axiomatic approach that does not require additional trainable parameters. Tang et al. (2022) have introduced a hierarchical graph learning model that offers a distinct approach to understanding brain region attributions. While our method is not hierarchical, it still effectively identifies region-specific attributions, offering a complementary perspective on connectomic features.

IG enabled us to systematically analyze attribution patterns across ROIs for each etiology. Figure 3 illustrates the mean node attributions for each etiology. As expected, the highest attributions are observed for nodes corresponding to their respective etiology classification, indicating that the model effectively identifies etiology-specific features.

To better understand the spatial distribution of the attributions, we mapped them to the corresponding ROIs of the ENA33 atlas, uncovering distinct etiology-specific patterns. For the control classification, regions across the brain contribute relatively evenly to the model’s predictions, reflecting a balanced distribution (Figure 4, A). This aligns with the expected variability within the control group, which comprises subjects from two datasets with potentially differing scanner characteristics.

In contrast, the attributions plots for PBs reveal a pronounced lateralization, with regions in the right hemisphere contributing more to the PB classification (Figure 5, A & B). The five most salient ENA33 ROIs for PB classification were exclusively right frontal and right parietal regions (Figure 5, C), which may reflect lateralized developmental features associated with preterm birth. This finding aligns with previous evidence suggesting that cerebral lateralization is impaired in prematurely-born individuals. Specifically, Kwon et al. (2015) demonstrated, using unbiased voxel-based measures of functional connectivity, that preterm neonates exhibit altered lateralization in left hemisphere language areas. The observed right-lateralized attribution pattern reflects the model’s sensitivity to input features that may capture aspects of these lateralization differences associated with preterm birth. Notably, while this lateralization is also present in other etiologies within the PB classification attribution plots, it is less pronounced (Figure 5, A).

For CHD, the attribution plots exhibit a less distinct pattern overall. However, examining the coronal plane renderings reveals heightened attribution in the left and right lateral temporal lobes compared to central regions of the brain (Figure 6, A & B). Interestingly, the five most salient ENA33 ROIs for CHD classification are all located in the right hemisphere, encompassing not only frontal and parietal regions but as suggested by Figure 6 (A) also two right temporal regions. The region with the highest attribution value for CHD classification in our data was the left Heschl’s gyrus. This finding resonates with previous evidence highlighting the importance of temporal regions in CHD. Jakab et al. (2019) demonstrated a strong correlation between the growth rate of the Heschl’s gyrus, anterior planum temporale, and language scores at 12 months of age in CHD patients. Their longitudinal study underscores the early role of left-dominant perisylvian regions in auditory and language development, suggesting that the attribution patterns observed here may capture subtle features related to these processes.

Lastly, for the SBA classification, attribution values show a trend within each hemisphere toward the right side, with the central part of the left hemisphere and the lateral part of the right hemisphere displaying the highest values (Figure 7, A & B). The five most salient ENA33 ROIs include not only temporal and parietal regions but also subcortical and occipital areas, marking a distinction from the other etiologies. Notably, no frontal regions were among the top five for SBA, emphasizing the unique role of subcortical regions in this etiology (Figure 7, C). This pattern may align with hydrocephalus-associated deformations, as hydrocephalus—a common feature of SBA—is known to alter the morphology of subcortical structures through increased intracranial pressure and disrupted cerebrospinal fluid flow (Paschereit et al., 2022).

When comparing the most salient regions across etiologies, the parietal inferior gyrus, Rolandic operculum, and frontal inferior gyrus consistently appear among the top regions for both PB and CHD classifications. These findings might reflect rapid changes in the white matter structure of these regions during development. The Rolandic operculum, a continuation of the pre- and postcentral gyri towards the lateral sulcus, is involved in emotion processing and functions as the sensory system for gustatory and visceral sensation (Sutoko et al., 2020). The parietal inferior lobule serves as an ’association area of association areas,’ integrating somatosensory, visual, and auditory information, critical for motor coordination and spatial awareness (Simpson & Fitch, 1988). Changes to these regions might indicate that regions undergoing most rapid development in the early neonatal period, such as regions associated with somatosensory functions, are most vulnerable for disorders. The frontal inferior gyrus, on the other hand, is involved in motor control and language processing, both of which are foundational for communication and higher-order cognitive functions (Liakakis et al., 2011). The attribution of these regions suggests their potential vulnerability to neurodevelopmental disruptions in PB and CHD etiologies. Notably, these regions do not appear among the most salient areas for the control classification, where occipital regions dominate instead (Figure 3C).

### 4.3 Limitations and future directions

Our study has the following limitations. First, the inherent differences in age ranges, scanner IDs, and mean sizes of the voxel connectomes between etiologies may introduce confounding factors. Although efforts were made to minimize these influences—such as analyzing embeddings with respect to potential confounders (see Supplementary Figure 1) and demonstrating through IG analysis that the model is not solely relying on confounding variables — these factors remain intrinsic to the datasets and may affect the generalizability of the findings. The use of data from only two sites further limits the diversity of the cohort, emphasizing the need for larger, multicenter datasets to validate these results across broader populations. Second, for graph learning purposes, studies would require more data to ensure robust generalization. We employed stratified sampling to address the dataset imbalance; however, expanding the dataset with additional samples, particularly from underrepresented etiologies, will be essential to enhance the reliability of these findings. Third, while the high spatial resolution of voxel-based parcellation is advantageous for capturing fine-grained connectomic details, it may also introduce increased noise, potentially overshadowing biologically meaningful differences and limiting the model’s interpretability. To mitigate this, we applied pruning by limiting each node to its top 256 edges which significantly improved learning outcomes. Exploring hybrid voxel-atlas parcellation strategies in future work could further reduce variability and improve interpretability. Additionally, we focused exclusively on GCNs, without exploring other GNN models, and we did not explicitly model edge features for performance reasons. Incorporating edge features could enhance model performance and enable edge-level attribution, providing additional insights into connectivity patterns. Future exploration of edge-level attributions within our framework could involve analyzing features associated with directed connectivity between nodes. Lastly, while this study focused on structural connectomes derived from diffusion MRI, incorporating multimodal data, such as functional MRI, could provide complementary information to further enhance classification accuracy. Future research should investigate the integration of multimodal data with graph learning methods to capture the full spectrum of connectomic alterations associated with neonatal etiologies.

By addressing these limitations and expanding the methodological framework, future studies can further refine the utility of graph-based machine learning for understanding neonatal connectomes and their relationship to neurodevelopmental outcomes.

### 4.4 Conclusion

This study demonstrates the potential of GCNs combined with voxel-based parcellation to classify neonatal connectomes by etiology, achieving an F1-score of 0.78. By leveraging the flexibility of voxel-based parcellation, we captured fine-grained connectomic differences, which improved classification performance compared to atlas-based methods. The integration of attribution analysis provided interpretable insights into etiology-specific patterns, highlighting the Rolandic operculum, inferior parietal lobule, and inferior frontal gyrus as regions of potentially particular importance for neurodevelopment in CHD and prematurity. This work marks a step toward atlas-independent connectome analysis, offering a versatile framework for studying diverse populations. By leveraging interpretable graph learning methods and validating them across diverse datasets, we move closer to enabling early identification of at-risk populations and paving the way for more tailored and effective interventions. Continued efforts in this direction will be crucial for bridging the gap between research insights and clinical applications.

## Acknowledgements

The authors want to first thank all families who participated in this research. In addition, we thank our contributing study group without whom the spina bifida research would not be possible. From the University Children’s Hospital this includes Barbara Casanova, Ruth Etter, Domenic Grisch, Mirjam Harm, Maya Horst Luethy, Jenny Kienzler, Niklaus Krayenbuehl, Markus A. Landolt, Andreas Meyer-Heim, Theres Moehrlen, Svea Muehlberg, Evelyne Riesen, Brigitte Seliner, Mithula Shellvarajah, Alexandra Wattinger and Noemi Zweifel. From the University Hospital Zurich, the study group consists of Lukas Kandler, Nele Struebing, Max Antonio Thomasius and Ladina Vonzun. Infrastructure support for the spina bifida research was provided by the Clinical Trial Center, University Hospital of Zurich. The authors also want to thank the The Swiss EPO Neuroprotection Trial Group: (alphabetical order by study site): Aarau: Neonatal Unit, Department of Pediatrics, Kantonsspital Aarau (Georg Zeilinger, MD; Sylviane Pasquier, MD), Department of Neuropediatrics (A. Capone). Basel: Universitätskinderklinik UKBB (Christoph Bührer, MD; René Glanzmann, MD; Sven Schulzke, MD), Department of Neuropediatric and Developmental Medicine (P. Weber). Chur: Abteilung für Neonatologie, Kantons-und Regionalspital (Brigitte Scharrer, MD; Walter Bär, MD), Department of Neuropediatrics (E. Keller, C. Killer). Geneva: Neonatology and Pediatric Intensive Care, Department of Pediatrics, University Hospital HCUG (Riccardo Pfister, MD; Krämer Karin, MD), Division of Development and Growth (P.S. Hüppi, C. Borradori-Tolsa). Zurich (Hans Ulrich Bucher, MD; Jean-Claude Fauchère, MD; Brigitte Koller, MSc.; Sven Wellmann, MD; Cornelia Hagmann, MD, PhD), University Children’s Hospital Zurich, Child Development Centre (B. Latal, G. Natalucci). The EPO study was supported by a grant received from the Swiss National Science Foundation (3200B0-108176) and was registered at clinicaltrials.gov (identifier: NCT 00313946).

Data were provided by the developing Human Connectome Project, KCL-Imperial-Oxford Consortium funded by the European Research Council under the European Union Seventh Framework Programme (FP/2007-2013) / ERC Grant Agreement no. [319456]. We are grateful to the families who generously supported this trial.

Also, the authors wish to acknowledge the Swiss National Supercomputing Center (CSCS) for providing computing infrastructure.

## Author Contribution

LG: conceptualization, data curation, formal analysis, investigation, methodology, model architecture, model training, project administration, software, validation, writing - original draft; AS: conceptualization, data curation, formal analysis, investigation, project administration, software, visualization, writing - original draft; SK: conceptualization, investigation, methodology, software, data analysis, visualization, validation, review and editing; AJ: data curation, funding acquisition, resources, supervision, review and editing; TH: funding acquisition, resources, supervision, review and editing.

## References

Agarap, A. F. (2018). Deep Learning using Rectified Linear Units (ReLU).

Andersson, J. L. R., & Sotiropoulos, S. N. (2016). An integrated approach to correction for off-resonance effects and subject movement in diffusion MR imaging. NeuroImage, 125, 1063–1078. 10.1016/j.neuroimage.2015.10.019

Avants, B. B., Tustison, N., & Johnson, H. (2014). Advanced Normalization Tools (ANTS) Release 2.x. https://brianavants.wordpress.com/2012/04/13/updated-ants-compile-instructions-april-12-2012/

Bastiani, M., Andersson, J. L. R., Cordero-Grande, L., Murgasova, M., Hutter, J., Price, A. N., Makropoulos, A., Fitzgibbon, S. P., Hughes, E., Rueckert, D., Victor, S., Rutherford, M., Edwards, A. D., Smith, S. M., Tournier, J.-D., Hajnal, J. V., Jbabdi, S., & Sotiropoulos, S. N. (2019). Automated processing pipeline for neonatal diffusion MRI in the developing Human Connectome Project. NeuroImage, 185, 750–763. 10.1016/j.neuroimage.2018.05.064

Batalle, D., Eixarch, E., Figueras, F., Muñoz-Moreno, E., Bargallo, N., Illa, M., Acosta-Rojas, R., Amat-Roldan, I., & Gratacos, E. (2012). Altered small-world topology of structural brain networks in infants with intrauterine growth restriction and its association with later neurodevelopmental outcome. NeuroImage, 60(2), 1352–1366. 10.1016/J.NEUROIMAGE.2012.01.059

Blesa, M., Serag, A., Wilkinson, A. G., Anblagan, D., Telford, E. J., Pataky, R., Sparrow, S. A., Macnaught, G., Semple, S. I., Bastin, M. E., & Boardman, J. P. (2016). Parcellation of the healthy neonatal brain into 107 Regions using atlas propagation through intermediate time points in childhood. Frontiers in Neuroscience, 10(MAY). 10.3389/fnins.2016.00220

Bonthrone, A. F., Chew, A., Bhroin, M. N., Rech, F. M., Kelly, C. J., Christiaens, D., Pietsch, M., Tournier, J.-D., Cordero-Grande, L., Price, A., Egloff, A., Hajnal, J. V, Pushparajah, K., Simpson, J., David Edwards, A., Rutherford, M. A., Nosarti, C., Batalle, D., & Counsell, S. J. (2022). Neonatal frontal-limbic connectivity is associated with externalizing behaviours in toddlers with Congenital Heart Disease. NeuroImage. Clinical, 36, 103153. 10.1016/j.nicl.2022.103153

Buckholtz, J. W., & Meyer-Lindenberg, A. (2012). Psychopathology and the Human Connectome: Toward a Transdiagnostic Model of Risk For Mental Illness. Neuron, 74(6), 990–1004. 10.1016/j.neuron.2012.06.002

Bullmore, E., & Sporns, O. (2009). Complex brain networks: graph theoretical analysis of structural and functional systems. Nature Reviews Neuroscience, 10(3), 186–198. 10.1038/nrn2575

Cai, T., Luo, S., Xu, K., He, D., Liu, T.-Y., & Wang, L. (2021). GraphNorm: A Principled Approach to Accelerating Graph Neural Network Training. In M. Meila & T. Zhang (Eds.), Proceedings of the 38th International Conference on Machine Learning (Vol. 139, pp. 1204–1215). PMLR. https://proceedings.mlr.press/v139/cai21e.html

Cao, M., Shu, N., Cao, Q., Wang, Y., & He, Y. (2014). Imaging Functional and Structural Brain Connectomics in Attention-Deficit/Hyperactivity Disorder. Molecular Neurobiology, 50(3), 1111–1123. 10.1007/s12035-014-8685-x

Cao, M., Yang, M., Qin, C., Zhu, X., Chen, Y., Wang, J., & Liu, T. (2021). Using DeepGCN to identify the autism spectrum disorder from multi-site resting-state data. Biomedical Signal Processing and Control, 70, 103015. 10.1016/j.bspc.2021.103015

Cordero-Grande, L., Christiaens, D., Hutter, J., Price, A. N., & Hajnal, J. V. (2019). Complex diffusion-weighted image estimation via matrix recovery under general noise models. NeuroImage, 200, 391–404. 10.1016/j.neuroimage.2019.06.039

Cordero-Grande, L., Hughes, E. J., Hutter, J., Price, A. N., & Hajnal, J. V. (2018). Three-dimensional motion corrected sensitivity encoding reconstruction for multi-shot multi-slice MRI: Application to neonatal brain imaging. Magnetic Resonance in Medicine, 79(3), 1365–1376. 10.1002/mrm.26796

Cui, H., Dai, W., Zhu, Y., Li, X., He, L., & Yang, C. (2021). BrainNNExplainer: An Interpretable Graph Neural Network Framework for Brain Network based Disease Analysis.

Cui, H., Dai, W., Zhu, Y., Li, X., He, L., & Yang, C. (2022). Interpretable Graph Neural Networks for Connectome-Based Brain Disorder Analysis.

Dadashkarimi, J., Karbasi, A., Liang, Q., Rosenblatt, M., Noble, S., Foster, M., Rodriguez, R., Adkinson, B., Ye, J., Sun, H., Camp, C., Farruggia, M., Tejavibulya, L., Dai, W., Jiang, R., Pollatou, A., & Scheinost, D. (2023). Cross Atlas Remapping via Optimal Transport (CAROT): Creating connectomes for different atlases when raw data is not available. Medical Image Analysis, 88, 102864. 10.1016/j.media.2023.102864

de Reus, M. A., & van den Heuvel, M. P. (2013). The parcellation-based connectome: Limitations and extensions. NeuroImage, 80, 397–404. 10.1016/j.neuroimage.2013.03.053

Desikan, R. S., Ségonne, F., Fischl, B., Quinn, B. T., Dickerson, B. C., Blacker, D., Buckner, R. L., Dale, A. M., Maguire, R. P., Hyman, B. T., Albert, M. S., & Killiany, R. J. (2006). An automated labeling system for subdividing the human cerebral cortex on MRI scans into gyral based regions of interest. NeuroImage, 31(3), 968–980. 10.1016/j.neuroimage.2006.01.021

Dhollander, T., Clemente, A., Singh, M., Boonstra, F., Civier, O., Duque, J. D., Egorova, N., Enticott, P., Fuelscher, I., Gajamange, S., Genc, S., Gottlieb, E., Hyde, C., Imms, P., Kelly, C., Kirkovski, M., Kolbe, S., Liang, X., Malhotra, A., … Caeyenberghs, K. (2021). Fixel-based Analysis of Diffusion MRI: Methods, Applications, Challenges and Opportunities. In NeuroImage (Vol. 241). Academic Press Inc. 10.1016/j.neuroimage.2021.118417

Dhollander, T., Mito, R., Raffelt, D., & Connelly, A. (2019, August). Improved white matter response function estimation for 3-tissue constrained spherical deconvolution. Proc. Intl. Soc. Mag. Reson. Med (Vol. 555, No. 10).

Dubois, J., Dehaene-Lambertz, G., Kulikova, S., Poupon, C., Hüppi, P. S., & Hertz-Pannier, L. (2014). The early development of brain white matter: A review of imaging studies in fetuses, newborns and infants. Neuroscience, 276, 48–71. 10.1016/j.neuroscience.2013.12.044

Edwards, A. D., Rueckert, D., Smith, S. M., Abo Seada, S., Alansary, A., Almalbis, J., Allsop, J., Andersson, J., Arichi, T., Arulkumaran, S., Bastiani, M., Batalle, D., Baxter, L., Bozek, J., Braithwaite, E., Brandon, J., Carney, O., Chew, A., Christiaens, D., … Hajnal, J. V. (2022). The Developing Human Connectome Project Neonatal Data Release. Frontiers in Neuroscience, 16. 10.3389/fnins.2022.886772

Fadnavis, S., Batson, J., & Garyfallidis, E. (2020). Patch2Self: Denoising Diffusion MRI with Self-Supervised Learning.

Feldmann, M., Bataillard, C., Ehrler, M., Ullrich, C., Knirsch, W., Gosteli-Peter, M. A., Held, U., & Latal, B. (2021). Cognitive and Executive Function in Congenital Heart Disease: A Meta-analysis. Pediatrics, 148(4). 10.1542/peds.2021-050875

Feldmann, M., Guo, T., Miller, S. P., Knirsch, W., Kottke, R., Hagmann, C., Latal, B., & Jakab, A. (2020). Delayed maturation of the structural brain connectome in neonates with congenital heart disease. Brain Communications, 2(2). 10.1093/braincomms/fcaa209

Fornito, A., Zalesky, A., Pantelis, C., & Bullmore, E. T. (2012). Schizophrenia, neuroimaging and connectomics. NeuroImage, 62(4), 2296–2314. 10.1016/j.neuroimage.2011.12.090

Galantucci, S., Agosta, F., Stefanova, E., Basaia, S., van den Heuvel, M. P., Stojković, T., Canu, E., Stanković, I., Spica, V., Copetti, M., Gagliardi, D., Kostić, V. S., & Filippi, M. (2017). Structural Brain Connectome and Cognitive Impairment in Parkinson Disease. Radiology, 283(2), 515–525. 10.1148/radiol.2016160274

He, K., Zhang, X., Ren, S., & Sun, J. (2016). Deep Residual Learning for Image Recognition. 2016 IEEE Conference on Computer Vision and Pattern Recognition (CVPR), 770–778. 10.1109/CVPR.2016.90

Hunt, D., Dighe, M., Gatenby, C., & Studholme, C. (2019). Challenges and Opportunities in Connectome Construction and Quantification in the Developing Human Fetal Brain. Topics in Magnetic Resonance Imaging : TMRI, 28(5), 265–273. 10.1097/RMR.0000000000000212

Jakab, A., Meuwly, E., Feldmann, M., Rhein, M. von, Kottke, R., O’Gorman Tuura, R., Latal, B., & Knirsch, W. (2019). Left temporal plane growth predicts language development in newborns with congenital heart disease. Brain, 142(5), 1270–1281. 10.1093/brain/awz067

Kellner, E., Dhital, B., Kiselev, V. G., & Reisert, M. (2016). Gibbs-ringing artifact removal based on local subvoxel-shifts. Magnetic Resonance in Medicine, 76(5), 1574–1581. 10.1002/mrm.26054

Kim, M., Yan, C., Yang, D., Liang, P., Kaufer, D. I., & Wu, G. (2021). Constructing Connectome Atlas by Graph Laplacian Learning. Neuroinformatics, 19(2), 233–249. 10.1007/s12021-020-09482-8

Kipf, T. N., & Welling, M. (2016). *Semi-Supervised Classification with Graph Convolutional Networks*.

Kollmorgen, S., Hahnloser, R. H. R., & Mante, V. (2020). Nearest neighbours reveal fast and slow components of motor learning. Nature, 577(7791), 526–530. 10.1038/s41586-019-1892-x

Kuklisova-Murgasova, M., Quaghebeur, G., Rutherford, M. A., Hajnal, J. V., & Schnabel, J. A. (2012). Reconstruction of fetal brain MRI with intensity matching and complete outlier removal. Medical Image Analysis, 16(8), 1550–1564. 10.1016/j.media.2012.07.004

Kwon, S. H., Scheinost, D., Lacadie, C., Sze, G., Schneider, K. C., Dai, F., Constable, R. T., & Ment, L. R. (2015). Adaptive mechanisms of developing brain: Cerebral lateralization in the prematurely-born. NeuroImage, 108, 144–150. 10.1016/j.neuroimage.2014.12.032

Li, X., Zhou, Y., Dvornek, N., Zhang, M., Gao, S., Zhuang, J., Scheinost, D., Staib, L. H., Ventola, P., & Duncan, J. S. (2021). BrainGNN: Interpretable Brain Graph Neural Network for fMRI Analysis. Medical Image Analysis, 74, 102233. 10.1016/j.media.2021.102233

Liakakis, G., Nickel, J., & Seitz, R. J. (2011). Diversity of the inferior frontal gyrus—A meta-analysis of neuroimaging studies. Behavioural Brain Research, 225(1), 341–347. 10.1016/J.BBR.2011.06.022

Loshchilov, I., & Hutter, F. (2017). Decoupled Weight Decay Regularization.

Maier-Hein, K. H., Neher, P. F., Houde, J.-C., Côté, M.-A., Garyfallidis, E., Zhong, J., Chamberland, M., Yeh, F.-C., Lin, Y.-C., Ji, Q., Reddick, W. E., Glass, J. O., Chen, D. Q., Feng, Y., Gao, C., Wu, Y., Ma, J., He, R., Li, Q., … Descoteaux, M. (2017). The challenge of mapping the human connectome based on diffusion tractography. Nature Communications, 8(1), 1349. 10.1038/s41467-017-01285-x

Makropoulos, A., Robinson, E. C., Schuh, A., Wright, R., Fitzgibbon, S., Bozek, J., Counsell, S. J., Steinweg, J., Vecchiato, K., Passerat-Palmbach, J., Lenz, G., Mortari, F., Tenev, T., Duff, E. P., Bastiani, M., Cordero-Grande, L., Hughes, E., Tusor, N., Tournier, J.-D., … Rueckert, D. (2018). The developing human connectome project: A minimal processing pipeline for neonatal cortical surface reconstruction. NeuroImage, 173, 88–112. 10.1016/j.neuroimage.2018.01.054

McCloskey, K., Taly, A., Monti, F., Brenner, M. P., & Colwell, L. J. (2019). Using attribution to decode binding mechanism in neural network models for chemistry. Proceedings of the National Academy of Sciences, 116(24), 11624–11629. 10.1073/pnas.1820657116

McCormick, M. C., Litt, J. S., Smith, V. C., & Zupancic, J. A. F. (2011). Prematurity: An Overview and Public Health Implications. Annual Review of Public Health, 32(1), 367–379. 10.1146/annurev-publhealth-090810-182459

McInnes, L., Healy, J., & Melville, J. (2018). UMAP: Uniform Manifold Approximation and Projection for Dimension Reduction.

Meuwly, E., Feldmann, M., Knirsch, W., von Rhein, M., Payette, K., Dave, H., Tuura, R. O. G., Kottke, R., Hagmann, C., Latal, B., Jakab, A., Liamlahi, R., Hackenberg, A., Kretschmar, O., Kellenberger, C., Bürki, C., & Weiss, M. (2019). Postoperative brain volumes are associated with one-year neurodevelopmental outcome in children with severe congenital heart disease. Scientific Reports, 9(1), 10885. 10.1038/s41598-019-47328-9

Moehrlen, U., Ochsenbein-Kölble, N., Mazzone, L., Kraehenmann, F., Hüsler, M., Casanova, B., Biro, P., Wille, D., Latal, B., Scheer, I., Bernet, V., Moehrlen, T., Held, L., Flake, A. W., Zimmermann, R., & Meuli, M. (2020). Benchmarking against the MOMS Trial: Zurich Results of Open Fetal Surgery for Spina Bifida. Fetal Diagnosis and Therapy, 47(2), 91–97. 10.1159/000500049

Möhrlen, U., Ochsenbein-Kölble, N., Mazzone, L., Kraehenmann, F., Hüsler, M., Casanova, B., Biro, P., Wille, D., Latal, B., Scheer, I., Bernet, V., Moehrlen, T., Held, L., Flake, A. W., Zimmermann, R., & Meuli, M. (2020). Benchmarking against the MOMS Trial: Zurich Results of Open Fetal Surgery for Spina Bifida. Fetal Diagnosis and Therapy, 47(2), 91–97. 10.1159/000500049

Murtagh, F. (1991). Multilayer perceptrons for classification and regression. Neurocomputing, 2(5–6), 183–197. 10.1016/0925-2312(91)90023-5

Nath, V., Schilling, K. G., Parvathaneni, P., Huo, Y., Blaber, J. A., Hainline, A. E., Barakovic, M., Romascano, D., Rafael-Patino, J., Frigo, M., Girard, G., Thiran, J.-P., Daducci, A., Rowe, M., Rodrigues, P., Prčkovska, V., Aydogan, D. B., Sun, W., Shi, Y., … Landman, B. A. (2020). Tractography reproducibility challenge with empirical data (TraCED): The 2017 ISMRM diffusion study group challenge. Journal of Magnetic Resonance Imaging : JMRI, 51(1), 234–249. 10.1002/jmri.26794

Ní Bhroin, M., Abo Seada, S., Bonthrone, A. F., Kelly, C. J., Christiaens, D., Schuh, A., Pietsch, M., Hutter, J., Tournier, J. D., Cordero-Grande, L., Rueckert, D., Hajnal, J. V., Pushparajah, K., Simpson, J., Edwards, A. D., Rutherford, M. A., Counsell, S. J., & Batalle, D. (2020). Reduced structural connectivity in cortico-striatal-thalamic network in neonates with congenital heart disease. NeuroImage: Clinical, 28. 10.1016/j.nicl.2020.102423

O’Gorman, R. L., Bucher, H. U., Held, U., Koller, B. M., Hüppi, P. S., & Hagmann, C. F. (2015). Tract-based spatial statistics to assess the neuroprotective effect of early erythropoietin on white matter development in preterm infants. Brain, 138(2), 388–397. 10.1093/brain/awu363

Papo, D., Buldú, J. M., Boccaletti, S., & Bullmore, E. T. (2014). Complex network theory and the brain. Philosophical Transactions of the Royal Society B: Biological Sciences, 369(1653), 20130520. 10.1098/rstb.2013.0520

Paschereit, F., Schindelmann, K. H., Hummel, M., Schneider, J., Stoltenburg-Didinger, G., & Kaindl, A. M. (2022). Cerebral Abnormalities in Spina Bifida: A Neuropathological Study. Pediatric and Developmental Pathology, 25(2), 107–123. 10.1177/10935266211040500

Payette, K., de Dumast, P., Kebiri, H., Ezhov, I., Paetzold, J. C., Shit, S., Iqbal, A., Khan, R., Kottke, R., Grehten, P., Ji, H., Lanczi, L., Nagy, M., Beresova, M., Nguyen, T. D., Natalucci, G., Karayannis, T., Menze, B., Cuadra, M. B., & Jakab, A. (2021). An automatic multi-tissue human fetal brain segmentation benchmark using the Fetal Tissue Annotation Dataset. Scientific Data, 8(1). 10.1038/s41597-021-00946-3

Rosenthal, G., Váša, F., Griffa, A., Hagmann, P., Amico, E., Goñi, J., Avidan, G., & Sporns, O. (2018). Mapping higher-order relations between brain structure and function with embedded vector representations of connectomes. Nature Communications, 9(1), 2178. 10.1038/s41467-018-04614-w

Rubinov, M., & Sporns, O. (2010). Complex network measures of brain connectivity: Uses and interpretations. NeuroImage, 52(3), 1059–1069. 10.1016/j.neuroimage.2009.10.003

Sanchez-Lengeling, B., Wei, J., Lee, B., Reif, E., Wang, P., Qian, W., McCloskey, K., Lucy, C., & Wiltschko, A. (2020). Evaluating Attribution for Graph Neural Networks. In H. Larochelle, M. Ranzato, R. Hadsell, M. F. Balcan, & H. Lin (Eds.), Advances in Neural Information Processing Systems (Vol. 33, pp. 5898–5910). Curran Associates, Inc. https://proceedings.neurips.cc/paper_files/paper/2020/file/417fbbf2e9d5a28a855a11894 b2e795a-Paper.pdf

Schneider, J., Mohr, N., Aliatakis, N., Seidel, U., John, R., Promnitz, G., Spors, B., & Kaindl, A. M. (2021). Brain malformations and cognitive performance in spina bifida. Developmental Medicine & Child Neurology, 63(3), 295–302. 10.1111/dmcn.14717

Simpson, J. A., & Fitch, W. (1988). Integrative functions of the cerebral cortex. Applied Neurophysiology, 109–116. 10.1016/B978-0-7236-0707-6.50013-2

Smith, L. N. (2017). Cyclical Learning Rates for Training Neural Networks. 2017 IEEE Winter Conference on Applications of Computer Vision (WACV), 464–472. 10.1109/WACV.2017.58

Smith, L. N., & Topin, N. (2017). Super-Convergence: Very Fast Training of Neural Networks Using Large Learning Rates.

Smith, R. E., Tournier, J.-D., Calamante, F., & Connelly, A. (2012). Anatomically-constrained tractography: Improved diffusion MRI streamlines tractography through effective use of anatomical information. NeuroImage, 62(3), 1924–1938. 10.1016/j.neuroimage.2012.06.005

Smith, R. E., Tournier, J.-D., Calamante, F., & Connelly, A. (2015a). SIFT2: Enabling dense quantitative assessment of brain white matter connectivity using streamlines tractography. NeuroImage, 119, 338–351. 10.1016/j.neuroimage.2015.06.092

Smith, R. E., Tournier, J.-D., Calamante, F., & Connelly, A. (2015b). The effects of SIFT on the reproducibility and biological accuracy of the structural connectome. NeuroImage, 104, 253–265. 10.1016/j.neuroimage.2014.10.004

Smith, S. M., Jenkinson, M., Woolrich, M. W., Beckmann, C. F., Behrens, T. E. J., Johansen-Berg, H., Bannister, P. R., De Luca, M., Drobnjak, I., Flitney, D. E., Niazy, R. K., Saunders, J., Vickers, J., Zhang, Y., De Stefano, N., Brady, J. M., & Matthews, P. M. (2004). Advances in functional and structural MR image analysis and implementation as FSL. NeuroImage, 23, S208–S219. 10.1016/j.neuroimage.2004.07.051

Speckert, A., Ji, H., Payette, K., Grehten, P., Kottke, R., Ackermann, S., Padden, B., Mazzone, L., Moehrlen, U., & Jakab, A. (2024). OSBA: An Open Neonatal Neuroimaging Atlas and Template for Spina Bifida Aperta. Data, 9(9), 107. 10.3390/data9090107

Speckert, A., Payette, K., Knirsch, W., von Rhein, M., Grehten, P., Kottke, R., Hagmann, C., Natalucci, G., Moehrlen, U., Mazzone, L., Ochsenbein-Kölble, N., Padden, B., Latal, B., & Jakab, A. (2024). Altered connectome topology in newborns at risk for cognitive developmental delay: a cross-etiologic study. 10.1101/2024.06.20.599853

Srivastava, N., Hinton, G., Krizhevsky, A., & Salakhutdinov, R. (2014). Dropout: A Simple Way to Prevent Neural Networks from Overfitting. In Journal of Machine Learning Research (Vol. 15).

Subaramya, S., Kokul, T., Nagulan, R., & Pinidiyaarachchi, U. A. J. (2022). Graph Neural Network based Alzheimer’s Disease Classification using Structural Brain Network. 2022 22nd International Conference on Advances in ICT for Emerging Regions (ICTer), 1–6. 10.1109/ICTer58063.2022.10024076

Sundararajan, M., Taly, A., & Yan, Q. (2017a). Axiomatic Attribution for Deep Networks. In D. Precup & Y. W. Teh (Eds.), Proceedings of the 34th International Conference on Machine Learning (Vol. 70, pp. 3319–3328). PMLR. https://proceedings.mlr.press/v70/sundararajan17a.html

Sundararajan, M., Taly, A., & Yan, Q. (2017b). Axiomatic Attribution for Deep Networks. In D. Precup & Y. W. Teh (Eds.), Proceedings of the 34th International Conference on Machine Learning (Vol. 70, pp. 3319–3328). PMLR. https://proceedings.mlr.press/v70/sundararajan17a.html

Sutoko, S., Atsumori, H., Obata, A., Funane, T., Kandori, A., Shimonaga, K., Hama, S., Yamawaki, S., & Tsuji, T. (2020). Lesions in the right Rolandic operculum are associated with self-rating affective and apathetic depressive symptoms for post-stroke patients. Scientific Reports, 10(1), 20264. 10.1038/s41598-020-77136-5

Tang, H., Guo, L., Fu, X., Qu, B., Ajilore, O., Wang, Y., Thompson, P. M., Huang, H., Leow, A. D., & Zhan, L. (2022). A Hierarchical Graph Learning Model for Brain Network Regression Analysis. Frontiers in Neuroscience, 16. 10.3389/fnins.2022.963082

Tournier, J.-D., Calamante, F., & Connelly, A. (2010). Improved probabilistic streamlines tractography by 2nd order integration over fibre orientation distributions. Proceedings of the International Society for Magnetic Resonance in Medicine, 1670.

Tournier, J.-D., Smith, R., Raffelt, D., Tabbara, R., Dhollander, T., Pietsch, M., Christiaens, D., Jeurissen, B., Yeh, C.-H., & Connelly, A. (2019). MRtrix3: A fast, flexible and open software framework for medical image processing and visualisation. NeuroImage, 202, 116137. 10.1016/j.neuroimage.2019.116137

Tsai, S.-Y. (2018). Reproducibility of structural brain connectivity and network metrics using probabilistic diffusion tractography. Scientific Reports, 8(1), 11562. 10.1038/s41598-018-29943-0

Tustison, N. J., Avants, B. B., Cook, P. A., Zheng, Y., Egan, A., Yushkevich, P. A., & Gee, J. C. (2010). N4ITK_Improved_N3_Bias_Correction. IEE Transaction on Medical Imaging, 29(6), 1310–1320.

van den Heuvel, M. P., Kersbergen, K. J., de Reus, M. A., Keunen, K., Kahn, R. S., Groenendaal, F., de Vries, L. S., & Benders, M. J. N. L. (2015). The Neonatal Connectome During Preterm Brain Development. Cerebral Cortex, 25(9), 3000–3013. 10.1093/cercor/bhu095

van den Heuvel, M. P., & Sporns, O. (2019). A cross-disorder connectome landscape of brain dysconnectivity. Nature Reviews Neuroscience, 20(7), 435–446. 10.1038/s41583-019-0177-6

Wu, D., Li, X., & Feng, J. (2021). Connectome-based individual prediction of cognitive behaviors via graph propagation network reveals directed brain network topology. Journal of Neural Engineering, 18(4), 0460a–3. 10.1088/1741-2552/ac0f4d

Xu, K., Hu, W., Leskovec, J., & Jegelka, S. (2019). *How Powerful are Graph Neural Networks?*

Xu, T., Nenning, K.-H., Schwartz, E., Hong, S.-J., Vogelstein, J. T., Goulas, A., Fair, D. A., Schroeder, C. E., Margulies, D. S., Smallwood, J., Milham, M. P., & Langs, G. (2020). Cross-species functional alignment reveals evolutionary hierarchy within the connectome. NeuroImage, 223, 117346. 10.1016/j.neuroimage.2020.117346

Yang, L., Cao, G., Zhang, S., Zhang, W., Sun, Y., Zhou, J., Zhong, T., Yuan, Y., Liu, T., Liu, T., Guo, L., Yu, Y., Jiang, X., Li, G., Han, J., & Zhang, T. (2025). Contrastive machine learning reveals species -shared and -specific brain functional architecture. Medical Image Analysis, 101, 103431. 10.1016/j.media.2024.103431

Yu, M., Sporns, O., & Saykin, A. J. (2021). The human connectome in Alzheimer disease — relationship to biomarkers and genetics. Nature Reviews Neurology, 17(9), 545–563. 10.1038/s41582-021-00529-1

Zalesky, A., Solowij, N., Yucel, M., Lubman, D. I., Takagi, M., Harding, I. H., Lorenzetti, V., Wang, R., Searle, K., Pantelis, C., & Seal, M. (2012). Effect of long-term cannabis use on axonal fibre connectivity. Brain, 135(7), 2245–2255. 10.1093/brain/aws136

Zhao, T., Xu, Y., & He, Y. (2019). Graph theoretical modeling of baby brain networks. NeuroImage, 185, 711–727. 10.1016/j.neuroimage.2018.06.038

