## Supplementary Material for "Atlas-independent brain connectome analysis at voxel-level granularity: graph convolutional networks for etiology classification in newborns"

#### 1. Figures

##### Supplementary Figure 1

*UMAP visualization of the connectome embedding colored by potential confounding variables*

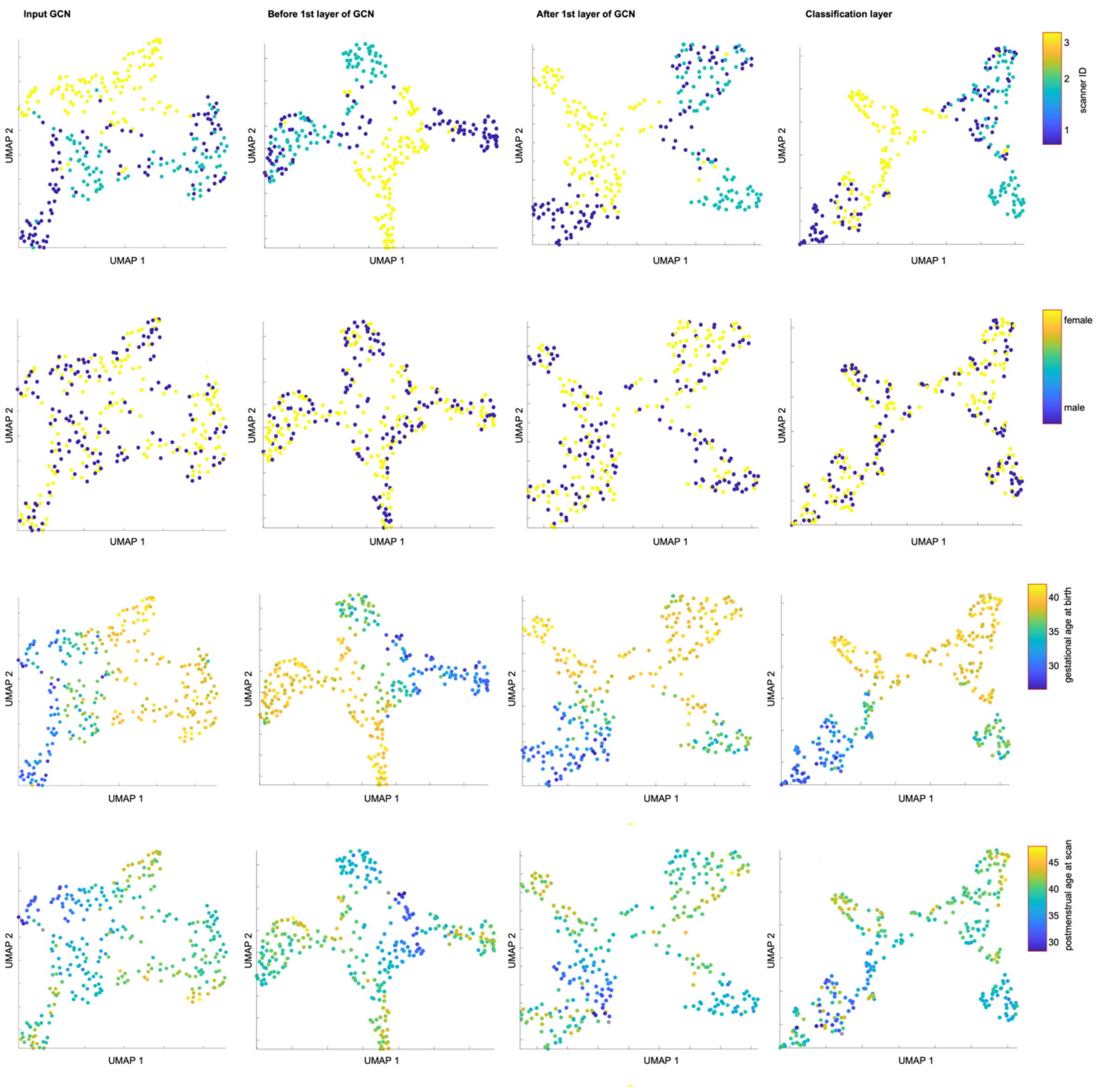


*Note.* Scanner ID 1 corresponds to the University Children's Hospital Zurich scanner before the software update, Scanner ID 2 to the same scanner after the update, and Scanner ID 3 to the dHCP scanner.

**Supplementary Figure 2**

*UMAP visualization of connectome embedding by etiology with individual subject trajectories highlighted*

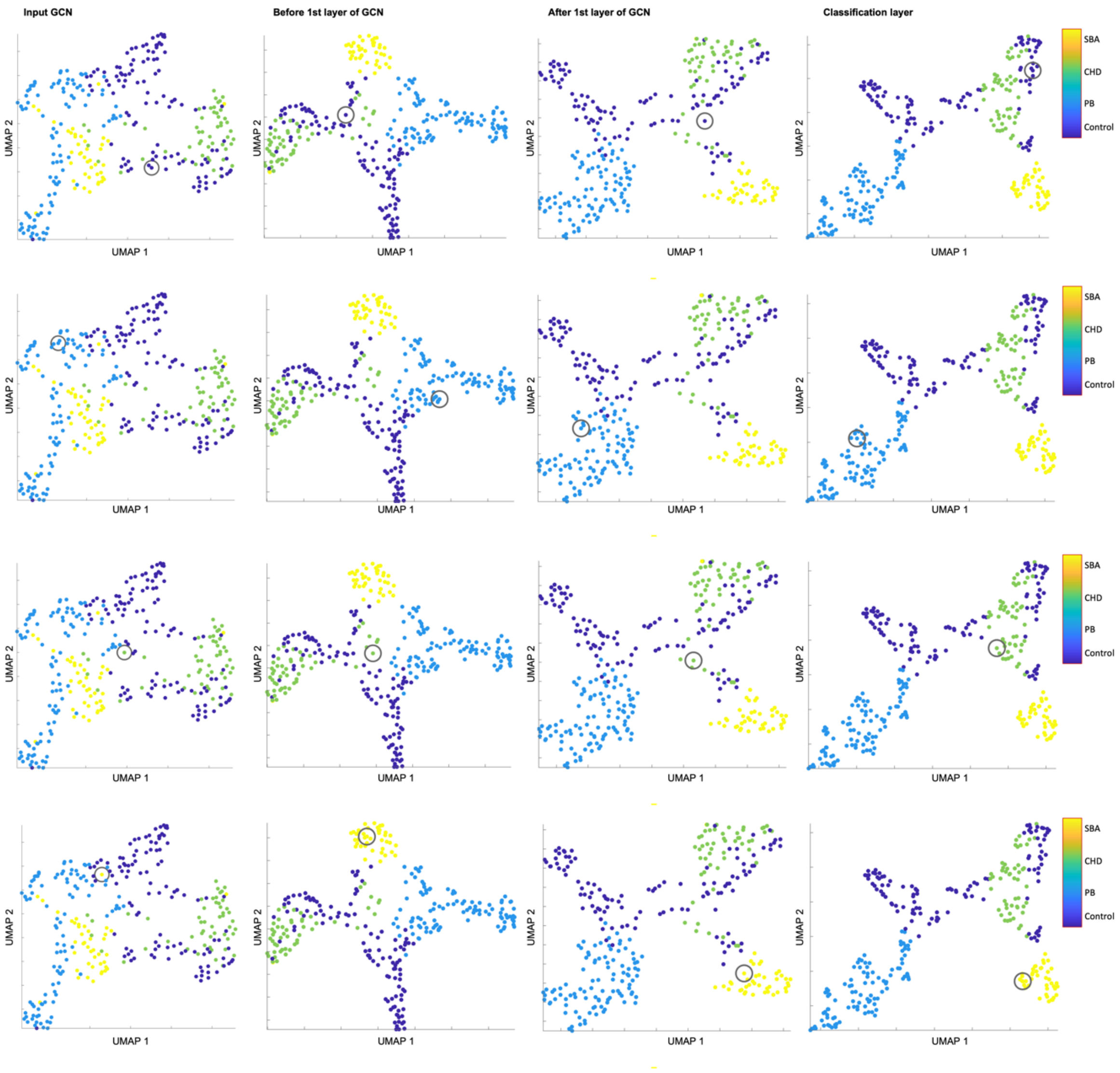


*Note.* The grey circle highlights one specific subject per etiology. In the first row, a Control subject is marked; in the second row, a PB (preterm-born) infant; in the third row, a CHD (congenital heart disease) infant; and in the fourth row, an SBA (spina bifida aperta) infant.

**Supplementary Figure 3**

*UMAP visualization of connectome embedding for Control and PB infants across scanner IDs*

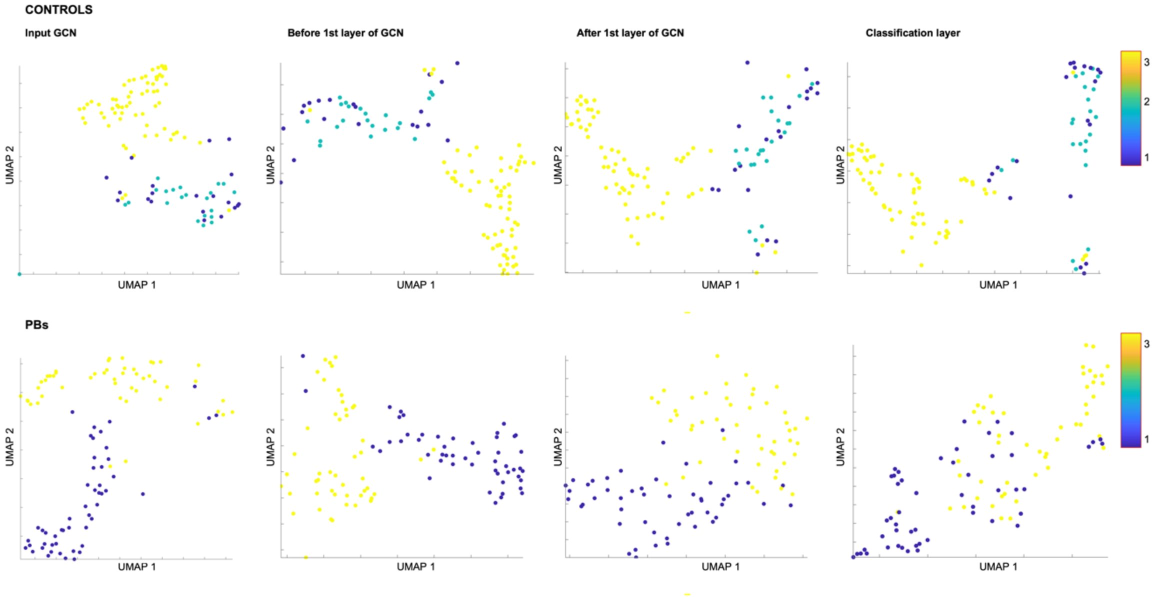


*Note.* In the top row the embeddings for the Control infants can found, in the bottom row the PB infants. Scanner ID 1 corresponds to the University Children's Hospital Zurich scanner before the software update, Scanner ID 2 to the same scanner after the update, and Scanner ID 3 to the dHCP scanner.

### 2. MATLAB Code

#### 2.1 Graph embeddings

##### 2.1.1 Filename: f_loadGraphEmbeddings.m

```
% (c) 2024, Sepp Kollmorgen, Anna Speckert

function [E, subjectIDs, foldIDs] = f_loadGraphEmbeddings(path, networkDepth, preFlag, dim)

   allFiles = dir(fullfile(path, sprintf('**/*_graph_h_%i%s.mat', networkDepth, preFlag)));

   E = NaN(numel(allFiles), dim);
   subjectIDs = cell(numel(allFiles), 1);
   foldIDs = cell(numel(allFiles), 1);

   for fid = 1:numel(allFiles)
      if mod(fid, 10) == 1
         fprintf('.');
      end
      filename = fullfile(path, allFiles(fid).name);
      L = load(filename);
      E(fid, :) = L.(sprintf('graph_h_%i%s', networkDepth, preFlag));
      ss = strsplit(filename, '_');
      if ~isempty(preFlag)
         subjectIDs{fid} = ss{end-4};
      else
         subjectIDs{fid} = ss{end-3};
      end
      foldIDs{fid} = ss{5};
   end

   fprintf('\n');

end
```

##### 2.1.2 Filename:  script_processGraphs.m

```
% (c) 2024, Sepp Kollmorgen, Anna Speckert

correctSlash = filesep;
folder = [correctSlash correctSlash fullfile('idnas37.d.uzh.ch', 'G_ADABD_Data$', 'Transfer', 'Connectomes', 'Processed')];
subFolder = fullfile('E_v4', 'matched_files');
nOutcomeClasses = 4;

% load graph embeddings for all subjects

% outcome prediction
[OP, subjectNames_op, foldNames] = f_loadGraphEmbeddings(fullfile(folder, subFolder), 2, '', nOutcomeClasses);

% graphEmbedding
[GE1pre, subjectNames_ge] = f_loadGraphEmbeddings(fullfile(folder, subFolder), 1, '_pre', 48);

% graphEmbedding
[GE1, subjectNames_ge1] = f_loadGraphEmbeddings(fullfile(folder, subFolder), 1, '', 48);

% graphEmbedding
[GE0, subjectNames_ge0] = f_loadGraphEmbeddings(fullfile(folder, subFolder), 0, '', 48);
```

```
assert(all(strcmp(subjectNames_op, subjectNames_ge)));

%isPostActivationFcn = [ones(size(E, 1), 1)*0; ones(size(E_, 1), 1)*1];

%% Get clinical information
clinicalFile = fullfile(folder, 'Auxillary', 'clinData_GNN_final.csv');
%clinicalFile = fullfile(folder, 'Auxillary', 'clinicalData.csv');
opts = detectImportOptions(clinicalFile);
opts = setvartype(opts,{'SubjectID', 'Etiology'},'char');
clinicalTable = readtable(clinicalFile, opts);

etiology = cell(numel(subjectNames_ge), 1);
sex = cell(numel(subjectNames_ge), 1);
pregnancyDuration = NaN(numel(subjectNames_ge), 1);
scanningAge = NaN(numel(subjectNames_ge), 1);
scannerUpgrade = NaN(numel(subjectNames_ge), 1);
cogComp_Score = NaN(numel(subjectNames_ge), 1);

for si = 1:numel(subjectNames_ge)

    ind = find(strcmp(clinicalTable.SubjectID, subjectNames_ge{si}));
    assert(numel(ind) == 1);
    etiology{si} = clinicalTable.Etiology{ind};
    sex{si} = clinicalTable.Sex{ind};
    pregnancyDuration(si) = clinicalTable.GA_birth(ind);
    scanningAge(si) = clinicalTable.PMA_scan(ind);
    scannerUpgrade(si) = clinicalTable.ScannerUpgrade(ind);
    cogComp_Score(si) = clinicalTable.CogComp_Score(ind);

end

dspace('E1', GE1, 'E1pre', GE1pre, 'E0', GE0, 'outcome_hat', OP, 'subjectID', subjectNames_ge,...
   'foldID', foldNames,...
   'Sex', sex, 'pregnancyDuration', pregnancyDuration, 'scanningAge', scanningAge,...
   'ScannerID', scannerUpgrade, 'cognitiveCompositeScore', cogComp_Score, 'Etiology', etiology);

% Reorder etiologies
dsource.Ldef.Etiology.Values = [0, 1, 2, 3];
dsource.Ldef.Etiology.ValueMeanings = {'Control', 'PB', 'CHD', 'SBA'};
newEtiology = NaN(dsource.N, 1);

newEtiology(dsource.L.Etiology == 1) = 2;
newEtiology(dsource.L.Etiology == 2) = 0;
newEtiology(dsource.L.Etiology == 3) = 1;
newEtiology(dsource.L.Etiology == 4) = 3;

dsource.L.Etiology = newEtiology;



E1 = dsource.F('E1').getMatrix(1:dsource.N);
dsource.L.nZeros1 = sum(abs(E1) < 1e-6, 2);

E0 = dsource.F('E0').getMatrix(1:dsource.N);
dsource.L.nZeros0 = sum(abs(E0) < 1e-6, 2);

dspaceApp.updateAll();


%% Compute embeddings
```

```
% parameters = [];
% parameters.empty = [];
% parameters.featureName = 'E0';
% parameters.useAllXf = false;
% parameters.pcaDimensions = 100;
% parameters.graph = '---';
% parameters.dimensions = 2;
% parameters.metric = 'euclidean';
% parameters.n_epochs = 1000;
% parameters.n_neighbours = 10;
% parameters.min_dist = 0.3;
% parameters.method = 'Java';
% parameters.Y0 = dspace.app.StringList({}, true(1, 0));
% parameters.zScore = true;
% parameters.supervisionLabel = '---';
% parameters.target_weight = 0.5;
% parameters.target_metric = 'categorical';
% parameters.isBackgroundComputation = false;
% parameters.variableName = 'UMAP_E0_eucl_';
% parameters.runAll = false;
%
% view = dspace.Dataview.getConfiguredDefaultView(dsource);
% [~, ~, results] = dspacefcns.essentials.features.umapOnCurrentSelection(view, parameters);
```

#### 2.2 Integrated Gradients Analysis

##### 2.2.1 Filename: f_loadSaliencies.m

```
% (c) 2024, Sepp Kollmorgen, Anna Speckert

function [S_features, S_nodes, nodeIndices] = f_loadSaliencies(path, validationFoldName, subjectId)

  filename = fullfile(path, sprintf('%s_integrated_gradients_%s.mat', validationFoldName, subjectId));

  L = load(filename);

  S_features = L.integrated_gradients_features;

  S_nodes = L.integrated_gradients_nodes;

  nodeIndices = (1:size(S_features, 1))';

end
```

##### 2.2.2 Filename: f_averageSalienciesPerArea.m

```
function f_averageSalienciesPerArea(dproject, binningLabel)

  for j = 1:numel(dproject.Sources.RAW)
    dsource = dproject.Sources.RAW{j};

    ls = {'atlasLabel', 'atlas__x', 'atlas__y', 'S_node_Control', 'S_node_PB', 'S_node_SBA', 'S_node_CHD',
'clinical__etiology'};
    %ls = dsource.getLabelNames();

    bins = unique(dsource.L.(binningLabel)(:));

    T = struct(); %'Size', [numel(bins), numel(ls)*4], 'VariableTypes', repmat({'single'}, 1, numel(ls)*4));
```

```
        for bi = 1:numel(bins)
            valid = dsource.L.(binningLabel) == bins(bi);
            for lsi = 1:numel(ls)
                X = dsource.L.(ls{lsi})(valid);
                X = X(:);
                if ismember(lsi, [4, 5, 6, 7])
                    T(bi).([ls{lsi} '_mean']) = mean(X, 'omitnan');
                    T(bi).([ls{lsi} '_std']) = std(X, 'omitnan');
                    T(bi).([ls{lsi} '_median']) = median(X, 'omitnan');
                    T(bi).([ls{lsi} '_iqr']) = iqr(X(~isnan(X)));
                else
                    T(bi).([ls{lsi}]) = mean(X, 'omitnan');
                end

            end
        end

        T = struct2table(T);

        newSource = dspace(T);
        newSource.Name = dsource.Name;
        for lsi = 1:numel(ls)
            if isfield(dsource.Ldef, ls{lsi})
                newSource.Ldef.(ls{lsi}) = dsource.Ldef.(ls{lsi});
            end
        end

        dproject.insertSource(newSource, 'ATLAS_AREA_AVG', true);
        fprintf('.');

    end

    fprintf('\n');

end
```

##### 2.2.3 Filename: f_assembleSubjectSaliencies.m

```
% (c) 2024, Sepp Kollmorgen, Anna Speckert

function dataset = f_assembleSubjectSaliencies(subjectId, validationFold, folder)


    % Find label file - thanks Anna!

    allFiles = dir(fullfile(folder, 'Auxillary', 'voxel2Label', sprintf('**/*%s_label_mapping.csv', subjectId)));

    assert(numel(allFiles) == 1);

    %fullfile(folder, 'E_v1', 'saliency', '**/*.mat'));
    subjectFilename = allFiles(1).name;
    labelFile = fullfile(folder, 'Auxillary', 'voxel2Label', subjectFilename);
    labelTable = readtable(labelFile);

    [S_features, S_nodes, nodeIndices] = f_loadSaliencies(fullfile(folder, 'E_v4', 'saliency'), validationFold, subjectId);

    S_features_Control = S_features(:, :, 1);
    S_features_PB = S_features(:, :, 2);
    S_features_CHD = S_features(:, :, 3);
    S_features_SBA = S_features(:, :, 4);
```

```
    S_node_Control = S_nodes(:, 1);
    S_node_PB = S_nodes(:, 2);
    S_node_CHD = S_nodes(:, 3);
    S_node_SBA = S_nodes(:, 4);



    % check consistency
    if max(labelTable.Voxel_Label) ~= numel(nodeIndices)
        fprintf('Label_mapping node count (%i) and embedding node count (%i) do not match.\n',...
            max(labelTable.Voxel_Label), max(nodeIndices));
    end

    atlasLabel = labelTable.ENA33_Label;

    dataset = dspace(S_features_Control, S_features_PB, S_features_CHD, S_features_SBA, ...
        S_node_Control, S_node_PB, S_node_CHD, S_node_SBA, nodeIndices, atlasLabel);

%     E_features = dataset.F('E').getMatrix(1:dataset.N);
%     dataset.L.saliency__mean = mean(E_features, 2);
%     dataset.L.saliency__max = max(E_features, [], 2);
%     dataset.L.saliency__min = min(E_features, [], 2);
%     dataset.L.saliency__var = var(E_features, [], 2);
%     dataset.L.saliency__range = max(abs(E_features), [], 2);
%     dataset.L.saliency__meanAbs = mean(abs(E_features), 2);
%     dataset.L.saliency__direction = mean(E_features(:, 4:end), 2);
%     dataset.L.saliency__position = mean(E_features(:, 1:3), 2);
%
%     dataset.L.saliency__direction_perc = invprctile(dataset.L.saliency__direction, dataset.L.saliency__direction);
%     dataset.L.saliency__position_perc = invprctile(dataset.L.saliency__position, dataset.L.saliency__position);
%     dataset.L.saliency__mean_perc = invprctile(dataset.L.saliency__mean, dataset.L.saliency__mean);
%     dataset.L.saliency__meanAbs_perc = invprctile(dataset.L.saliency__meanAbs, dataset.L.saliency__meanAbs);


    prcTransform = {'S_node_Control', 'S_node_PB', 'S_node_CHD', 'S_node_SBA'};

    for k = 1:numel(prcTransform)
        dataset.L.([prcTransform{k} '_perc']) = invprctile(dataset.L.(prcTransform{k}), dataset.L.(prcTransform{k}));
    end

    dataset.Name = subjectFilename(1:end-17);

end
```

##### 2.2.4 Filename: script_processSalienciesAcrossSubjects.m

```
correctSlash = filesep;

folder = [correctSlash correctSlash fullfile('idnas37.d.uzh.ch', 'G_ADABD_Data$', 'Transfer', 'Connectomes', 'Processed')];

atlasFile = fullfile(folder, 'Auxillary', 'Atlas_averageBrain_ realWorldCoord.csv');
atlasTable = readtable(atlasFile);

clinicalFile = fullfile(folder, 'Auxillary', 'clinData_GNN_final.csv');
opts = detectImportOptions(clinicalFile);
opts = setvartype(opts,{'SubjectID'},'char');
clinicalTable = readtable(clinicalFile, opts);

allFiles = dir(fullfile(folder, 'E_v4', 'saliency', '**/*.mat'));

allDatasets = cell(1, numel(allFiles));
for fid = 1:numel(allFiles)
    fprintf('%s...\n', allFiles(fid).name);
```

```
    ss = strsplit(allFiles(fid).name, '_integrated_gradients_');
    subjectId = ss{end}(1:end-4);
    validationFold = ss{1};

    dataset = f_assembleSubjectSaliencies(subjectId, validationFold, folder);

    % Add atlas information

    aDef = dspace_data.PropertyDefinition(atlasTable.Label, atlasTable.ROI', '', '', []);
    dataset.Ldef.atlasLabel = aDef;
    dataset.Ldef.atlasLabel.valueTable = atlasTable;

    valid = dataset.L.atlasLabel <= 94;

    dataset.L.atlas__x = NaN(dataset.N, 1);
    dataset.L.atlas__y = NaN(dataset.N, 1);
    dataset.L.atlas__z = NaN(dataset.N, 1);
    dataset.L.atlas__mass = NaN(dataset.N, 1);
    dataset.L.atlas__lobe = NaN(dataset.N, 1);

    dataset.L.atlas__x(valid) = atlasTable.x(dataset.L.atlasLabel(valid));
    dataset.L.atlas__y(valid) = atlasTable.y(dataset.L.atlasLabel(valid));
    dataset.L.atlas__z(valid) = atlasTable.z(dataset.L.atlasLabel(valid));
    dataset.L.atlas__mass(valid) = atlasTable.Mass(dataset.L.atlasLabel(valid));
    dataset.L.atlas__lobe(valid) = atlasTable.ModuleID(dataset.L.atlasLabel(valid));
    lobeDef = dspace_data.PropertyDefinition(1:6, {'Frontal', 'Parietal', 'Temporal', 'Occipital',...
        'Limbic and Insula', 'Subcortical and cerebellum'}, '', '', []);
    dataset.Ldef.atlas__lobe = lobeDef;

    ind = find(strcmp(clinicalTable.SubjectID, subjectId));
    assert(numel(ind) == 1);

    subTable = table();
    subTable.clinical__ageAtBirth_weeks = NaN(dataset.N, 1);

    subTable.clinical__etiology(:) = {clinicalTable.Etiology{ind}};
    subTable.clinical__ageAtBirth_weeks(:) = clinicalTable.GA_birth(ind);
    subTable.clinical__ageAtScan_weeks(:) = clinicalTable.PMA_scan(ind);
    subTable.clinical__sex(:) = {clinicalTable.Sex{ind}};
    subTable.clinical__scannerUpgrade(:) = clinicalTable.ScannerUpgrade(ind);
    subTable.clinical__CogComp_Score(:) = clinicalTable.CogComp_Score(ind);

    dsubTable = dspace(subTable);
    dsubTable.L.idx = [];
    dataset.L = [dataset.L dsubTable.L];
    dataset.Ldef.clinical__etiology = dsubTable.Ldef.clinical__etiology;
    dataset.Ldef.clinical__sex = dsubTable.Ldef.clinical__sex;

    % + Outcome

    allDatasets{fid} = dataset;

end

dproject = dspace_data.DataCollection({allDatasets}, {'RAW'}, 'NodeSaliencies');
dspace(dproject);
f_averageSalienciesPerArea(dproject, 'atlasLabel');

%% Reorder etiologies (after merging datasets)
dsource.Ldef.Etiology = dsource.Ldef.clinical__etiology;
dsource.Ldef.Etiology.Values = [0, 1, 2, 3];
dsource.Ldef.Etiology.ValueMeanings = {'Control', 'PB', 'CHD', 'SBA'};
```

```
newEtiology = NaN(dsource.N, 1);

newEtiology(dsource.L.clinical__etiology == 3) = 0;
newEtiology(dsource.L.clinical__etiology == 4) = 1;
newEtiology(dsource.L.clinical__etiology == 2) = 2;
newEtiology(dsource.L.clinical__etiology == 1) = 3;

dsource.L.Etiology = newEtiology;

%% Compute percentiles
ls = {'S_node_Control_mean', 'S_node_PB_mean', 'S_node_SBA_mean', 'S_node_CHD_mean'};

for lsi = 1:numel(ls)
    dsource.L.([ls{lsi} '_invperc']) = invprctile(dsource.L.(ls{lsi}), dsource.L.(ls{lsi}));
end
```
